# Genetic and phosphoproteomic basis of LysM-mediated immune signaling in *Marchantia polymorpha* highlights conserved elements and new aspect of pattern-triggered immunity in land plants

**DOI:** 10.1101/2022.12.28.521631

**Authors:** Izumi Yotsui, Hidenori Matsui, Shingo Miyauchi, Hidekazu Iwakawa, Katharina Melkonian, Titus Schlüter, Selena Gimenez Ibanez, Takehiko Kanazawa, Yuko Nomura, Sara Christina Stolze, Hyung-Woo Jeon, Shigeo Sugano, Makoto Shirakawa, Ryuichi Nishihama, Yasunori Ichihashi, Ken Shirasu, Takashi Ueda, Takayuki Kohchi, Hirofumi Nakagami

**Affiliations:** RIKEN Center for Sustainable Resource Science, Yokohama, Kanagawa, 230-0045 Japan; Department of BioScience, Tokyo University of Agriculture, Setagaya, Tokyo, 156-8502 Japan; Graduate School of Environmental and Life Sciences, Okayama University, Okayama, 700-8530 Japan; Max Planck Institute for Plant Breeding Research, 50829 Cologne, Germany; Okinawa Institute of Science and Technology Graduate University, Onna, Okinawa, 904-0495 Japan; School of Biological Science and Technology, College of Science and Engineering, Kanazawa University, Kakuma-machi, Kanazawa, Ishikawa, 920-1192 Japan; Department of Plant Molecular Genetics, Centro Nacional de Biotecnología, Consejo Superior de Investigaciones Científicas (CNB-CSIC), 28049 Madrid, Spain; Division of Cellular Dynamics, National Institute for Basic Biology, Nishigonaka 38, Myodaiji, Okazaki, Aichi, 444-8585 Japan; The Department of Basic Biology, SOKENDAI (The Graduate University for Advanced Studies), Nishigonaka 38, Myodaiji, Okazaki, Aichi, 444-8585 Japan; Graduate School of Biostudies, Kyoto University, Kyoto, 606-8502 Japan; Division of Biological Science, Graduate School of Science and Technology, Nara Institute of Science and Technology (NAIST), Ikoma, Nara, 630-0192 Japan; Department of Applied Biological Science, Faculty of Science and Technology, Tokyo University of Science, Noda, Chiba, 278-8510, Japan; RIKEN BioResource Research Center, Tsukuba, Ibaraki, 305-0074 Japan

## Abstract

Pattern-recognition receptor (PRR)-triggered immunity (PTI) wards off a wide range of pathogenic microbes, playing a pivotal role in plant immunity. The model liverwort *Marchantia polymorpha* is emerging as a popular model for investigating the evolution of plant-microbe interactions. *M. polymorpha* triggers defense-related gene expression upon sensing components of bacterial and fungal extracts, suggesting the existence of PTI in this plant model. However, the molecular components of the putative PTI in *M. polymorpha* have not yet been described. We show that, in *M. polymorpha*, which has four LysM receptor homologs, lysin motif (LysM) receptor-like kinase (LYK) MpLYK1 and LYK-related (LYR) MpLYR are required for sensing chitin and peptidoglycan fragments, triggering a series of characteristic immune responses. Comprehensive phosphoproteomic analysis of *M. polymorpha* in response to chitin treatment identified regulatory proteins that potentially shape LysM-mediated PTI. The identified proteins included homologs of well-described PTI components in angiosperms as well as proteins whose roles in PTI are not yet determined, including the blue-light receptor phototropin MpPHOT. We revealed that MpPHOT is required for a negative feedback of defense-related gene expression during PTI. Taken together, this study outlines the basic framework of LysM-mediated PTI in *M. polymorpha* and demonstrates the utility of *M. polymorpha* as a plant model for discovering novel or fundamental molecular mechanisms underlying PRR-triggered immune signaling in plants.

## Introduction

In angiosperms, cell-surface localized pattern-recognition receptors (PRRs) recognizing microbe-derived or plant-endogenous molecules play central roles in various plant-microbe interactions, which can be beneficial, neutral, or detrimental (Zipfel and Oldroyd, 2017). PRRs recognize slowly evolving microbe-associated molecular patterns (MAMPs) or symbiotic signals such as rhizobial nodulation (Nod) factors and mycorrhizal (Myc) factors. PRRs are transmembrane kinases or membrane-associated proteins that function together with kinases, triggering phosphorylation and/or interaction-dependent signaling to activate pattern-triggered immunity (PTI) or to initiate symbiosis (Zipfel and Oldroyd, 2017). Activation of PRRs typically induces a series of characteristic responses, including reactive oxygen species (ROS) production, MAP kinase (MAPK) activation, calcium influx, callose deposition, defense-related gene expression, and growth inhibition (Boller and Felix, 2009).

Well-studied bacterial MAMP receptors such as *Arabidopsis thaliana* AtFLS2 and AtEFR belong to the subfamily XII of leucine-rich repeat receptor-like kinases (LRR-RLKs). AtFLS2 and AtEFR recognize peptide fragments derived from bacterial flagellin and elongation factor Tu (EF-Tu), respectively (Chinchilla et al., 2006; Zipfel et al., 2006). These LRR-RLK subfamily XII MAMP receptors typically require SERK co-receptors belonging to the subfamily II of LRR-RLKs, for downstream signaling (Chinchilla et al., 2007; Roux et al., 2011). In *A. thaliana*, AtBAK1/AtSERK3 functions as a co-receptor for AtFLS2 and AtEFR as well as the brassinosteroid receptor AtBRI1, and thereby regulates not only PTI but also plant growth and development (Li et al., 2002).

N-acetylglucosamine (GlcNAc) derivatives from bacteria, fungi, and oomycetes, which can be either MAMPs or symbiotic signals, are perceived by lysin motif (LysM)-domain-containing receptors (Zipfel et al., 2017). As major cell wall components, bacterial peptidoglycans (PGN) and fungal chitin oligosaccharides are recognized as MAMPs by LysM receptor-like kinase (LYK), LYK-related (LYR), or LysM receptor-like protein (LYP) to activate PTI (Gust et al., 2012). In *A. thaliana*, chitin is perceived by LYR AtLYK5 and LYK AtCERK1, and PGN is perceived by LYP AtLYM1/3 and LYK AtCERK1 (Cao et al., 2014; Miya et al., 2007; Willmann et al., 2011). In rice, different combinations of LysM receptors perceive chitin and PGN (Kaku et al., 2006; Liu et al., 2012; Shimizu et al., 2010). In either case, LYK CERK1 most likely functions as a co-receptor as do the subfamily II LRR-RLK SERKs for the subfamily XII LRR-RLKs. Importantly, Nod and Myc factors are also perceived by LysM proteins to ensure beneficial symbiotic interactions. In *Lotus japonicus* and *Medicago truncatula*, different CERK1 homologs, most likely originating from gene duplications, function independently for PTI and symbiosis (Bozsoki et al., 2017). Intriguingly, however, rice LYK OsCERK1 is involved in both PTI and arbuscular mycorrhizal (AM) symbiosis (Carotenuto et al., 2017; Miyata et al., 2014; Zhang et al., 2015).

Considering the importance of PRRs in angiosperms for communication with various microbes, it is possible that acquisition and diversification of PRRs and their downstream signaling networks played a key role during plant terrestrialization and evolution. Homologs of characterized PRRs can be found in genomes of bryophytes and the charophyte alga *Chara braunii* (Bowman et al., 2017; Li et al., 2020; Nishiyama et al., 2018; Rensing et al., 2008; Zhang et al., 2020). The moss *Physcomitrium patens* has been shown to sense chitin and PGN fragments in a LYK PpCERK1-dependent manner (Bressendorff et al., 2016). The liverwort *M. polymorpha* is able to sense bacterial and fungal extracts, although the molecular components and mechanisms of this sensing are not yet described (Gimenez-Ibanez et al., 2019; Redkar et al., 2022). Here, we identify LysM receptors responsible for chitin- and PGN-induced responses in *M. polymorpha*, and provide evidence that the LysM receptor contributes to resistance against bacterial pathogens in *M. polymorpha*. Furthermore, we characterize the LysM-mediated signaling pathway in *M. polymorpha* by phosphoproteomics.

## Results

### *Marchantia polymorpha* recognizes chitin and PGN to induce immune responses

A rapid and transient burst of reactive oxygen species (ROS) is a hallmark of PRR activation upon MAMP perception in angiosperms. To investigate the conservation of MAMP recognition, we treated the wild-type *M. polymorpha* strains Tak-1 (male) and Tak-2 (female) with the known MAMPs, flg22, elf18, chitin, PGN, and lipopolysaccharide (LPS), which trigger ROS burst in *A. thaliana*. Chitin and PGN fragments, but not the other MAMPs, induced ROS burst in *M. polymorpha* (Figure 1a and 1b). In angiosperms and moss, LysM receptors are indispensable for sensing chitin and PGN. As in the case of angiosperms, long-chain chitin oligosaccharides induced stronger ROS burst (Figure 1c). Chitin treatment further induced MAP kinase (MAPK) activation, which was monitored by the use of an anti-p44/42-ERK antibody (a-pTEpY), as well as *WRKY* gene expression (Figure 1d and Supplementary Figure S3). These observations imply that LysM receptor-mediated MAMP perception and signaling mechanisms are conserved in *M. polymorpha*. We then investigated the chitin-induced transcriptional response in *M. polymorpha* Tak-2. Significant transcriptional reprogramming was observed 1 and 3 hours after chitin treatment (Figure 1e), and the transient nature of this response was similar to the chitin response in *A. thaliana* (Supplementary Figure S2a). Gene Ontology (GO) analysis of differentially expressed genes (DEGs, |Log_2_FC| > 2, adjusted p < 0.05) revealed that chitin treatment significantly and primarily induces expression of defense-related genes in *M. polymorpha* (Figure 1f) as in *A. thaliana* (Supplementary Figure S2b). Collectively, these results suggest the existence of a LysM-mediated immune signaling pathway in *M. polymorpha*.

**Figure 1.**
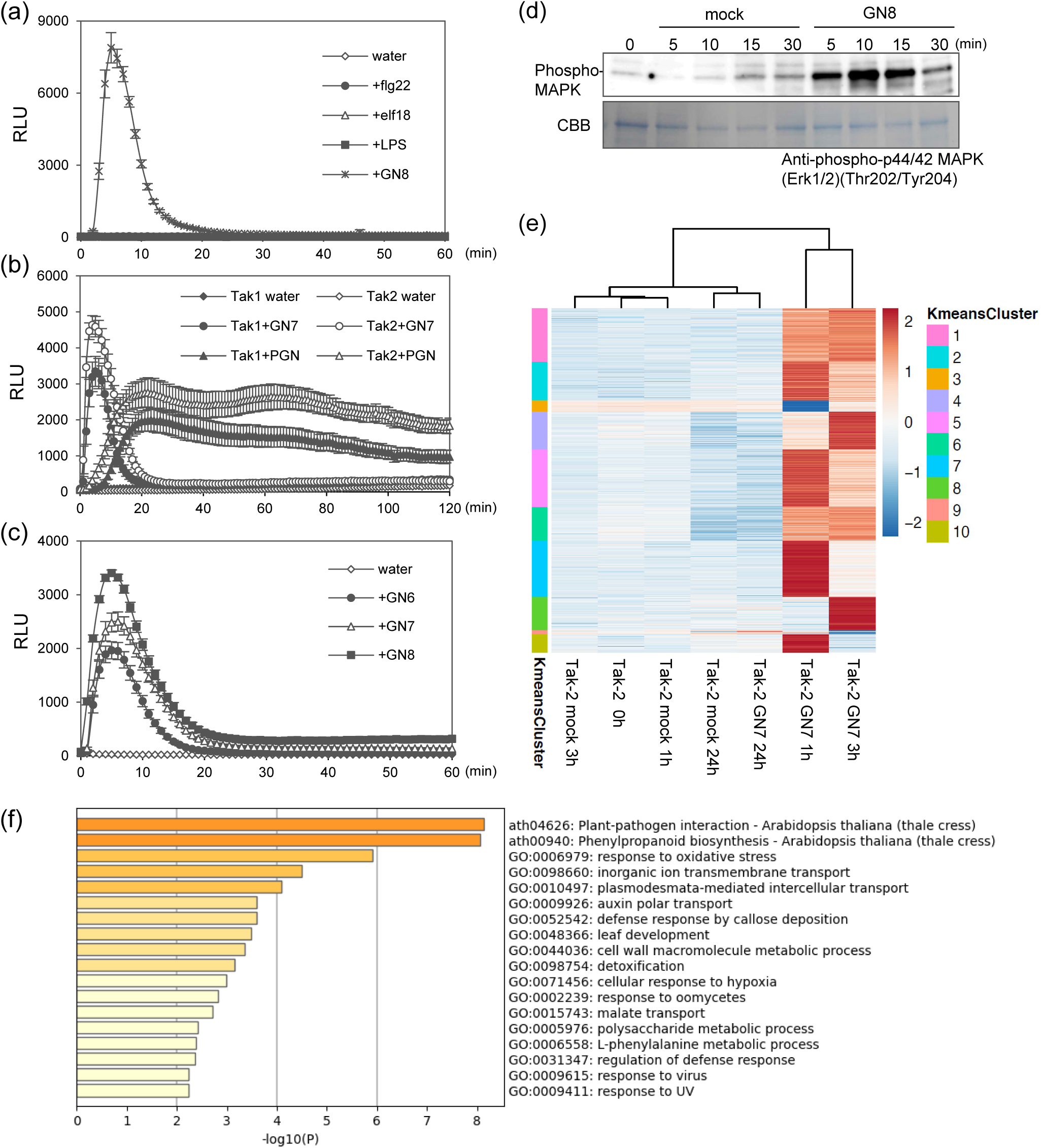
MAMP responses in *M. polymorpha*. (a-c) MAMP-induced reactive oxygen species (ROS) production in Tak-1 (male wild-type) and Tak-2 (female wild-type). ROS production after MAMP treatment was measured by chemiluminescence mediated by L-012 in 6-day-old gemmalings. (a) Wild-type gemmalings were treated with 1 μM flg22, 1 μM elf18, 100 μg/ml lipopolysaccharide (LPS) derived from *Pseudomonas aeruginosa*, or 1 μg/ml N-acetylchitooctaose (GN8) with HRP (horseradish peroxidase). (b) Wild-type gemmalings were treated with 1 μM N-acetylchitoheptaose (GN7) or 500 μg/ml peptidoglycan (PGN) derived from *Bacillus subtilis*. (c) Wild-type gemmalings were treated with 1 μM N-acetylchitohexaose (GN6), GN7, or GN8. The values represent the average and standard errors of four replicates. (d) GN8-induced MAPK activation in Tak-1. Tak-1 gemmalings were treated with 1 μg/ml GN8 for the indicated times. Activated MAPKs were detected by immunoblotting using anti-p44/42 MAPK antibody. (e) Clusters of *M. polymorpha* DEGs. Significantly differentially expressed genes with over ±2 log2 fold changes (FDR adjusted p < 0.05) were grouped based on K-means clustering. K-means cluster ID is shown on the left bar. Log2 read count of genes was normalized into the range of ±2. See Supplementary Table S7. (f) Enriched GO terms in the *M. polymorpha* DEGs (Fig. 1e). See Supplementary Table S8.

### MpLYK1 and MpLYR are responsible for chitin and PGN responses in *M. polymorpha*

LysM receptors can be roughly classified into LYK (LysM-RLK, LysM receptor-like kinase), LYR (LYK-related, LysM-RLK without classically conserved kinase domain), and LYP (LysM receptor-like protein, membrane-anchored LysM protein) (Gust et al., 2012; Tanaka et al., 2013). BLAST search identified four LysM receptor homologs, two LYKs, one LYR, and one LYP, in the genome of *M. polymorpha*. Phylogenetic analysis of the LysM domains of LysM receptor homologs in selected plant species, covering hornworts, liverwort, mosses, lycophyte, angiosperms, and streptophyte algae (Supplementary Table S1 and S2), revealed that the LysM domains of embryophytes form four major clades, LYKa, LYKb, LYR, and LYP (Figure 2a and Supplementary Figure S4). We found single *M. polymorpha* genes in each of the four clades, MpLYK1 (LYKa), MpLYK2 (LYKb), MpLYR, and MpLYP, which is in clear contrast to the moss *P. patens* that lacks *LYP* and *LYKb* but has instead evolved to harbor additional copies of *LYKa* and *LYR* (Figure 2a, Supplementary Table S1 and Figure S1). This suggests that *M. polymorpha* is a useful bryophyte model for comprehensive study of the function and molecular evolution of LysM receptors. MpLYK1 is orthologous to AtCERK1, MpLYK2 is orthologous to AtLYK3, and MpLYR is orthologous to AtLYK4 and AtLYK5 (Figure 2a). LysM domains from Charophyceae, *Chara braunii* and *Nitella mirabilis*, formed another independent clade (Figure 2a and Supplementary Table S1). It is possible that embryophyte LysM receptors were derived from a LysM receptor in the algal ancestor of embryophytes.

**Figure 2.**
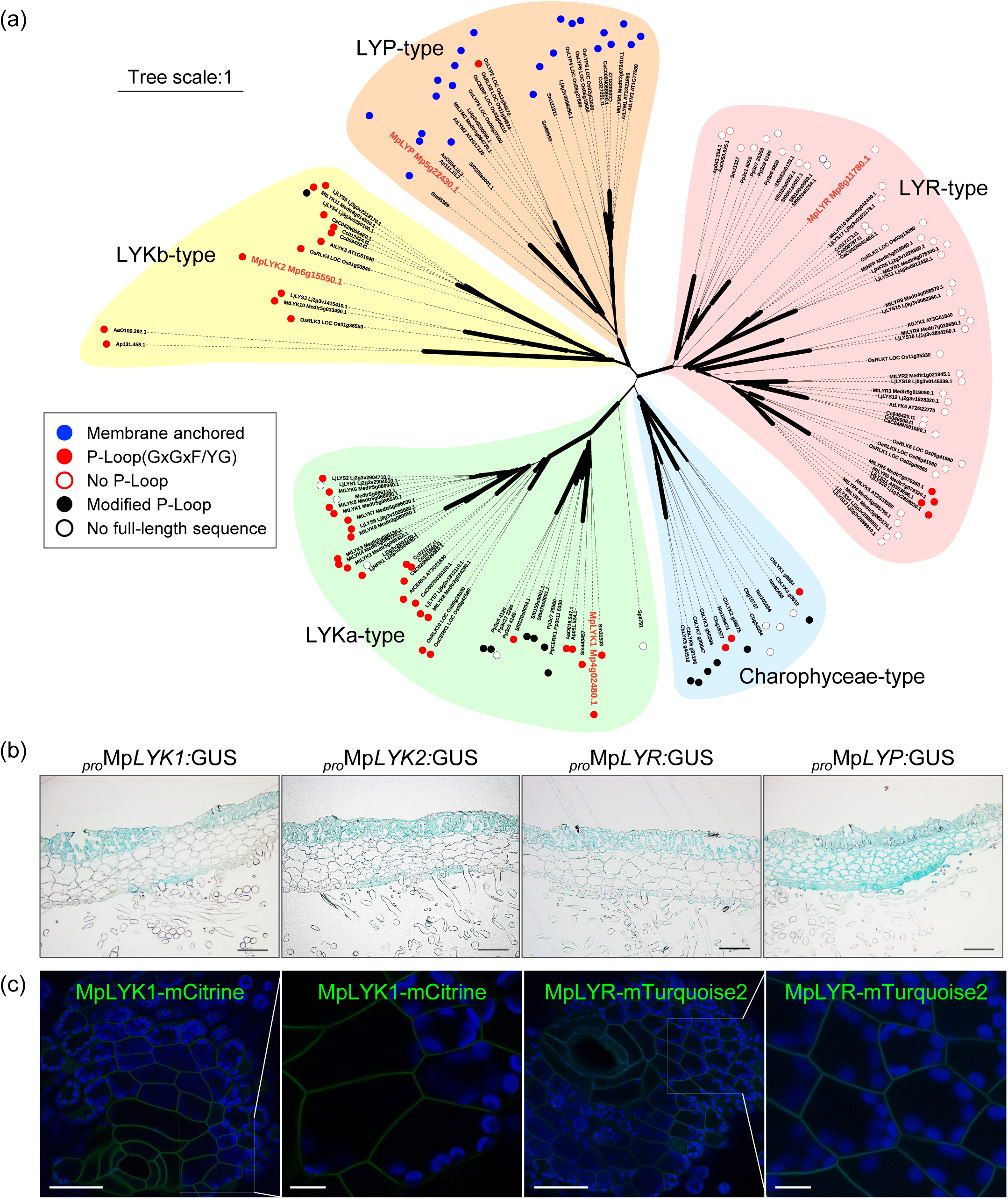
LysM receptor homologs in *M. polymorpha*. (a) Unrooted phylogenetic tree of LysM proteins in plants. Amino acid sequences of ectodomain including LysM1, LysM2, and LysM3 domain were used for drawing the tree. A graphical view of the tree has been generated using iTOL. Width of branches denote bootstrap support based on 1000 repetitions. Major subgroups were designated as LYKa, LYKb, LYR, LYP, and Charophyceae type. The proteins containing the basic P-Loop (GxGxF/YG) or no P-Loop are shown by red circles or red flames, respectively. The proteins containing modified P-Loops (See Supplementary Table S2) or no full-length sequences in the databases are shown by black circles or black flames, respectively. The membrane anchored-type proteins are shown by blue circles. *M. polymorpha* proteins are highlighted in red letters. Abbreviations: At, *Arabidopsis thaliana*; Mt, *Medicago truncatula*; Lj, *Lotus japonicus*; Os, *Oryza sativa*; Ca, *Cuscuta australis*; Cc, *Cuscuta campestris*; Sm, *Selaginella moellendorffii*; Sf, *Sphagnum fallax*; Pp, *Physcomitrium patens*; AaO, *Anthoceros agrestis* [Oxford]; Ap, *Anthoceros punctatus*; Mp, *Marchantia polymorpha*; Nm, *Nitella mirabilis*; Cb, *Chara braunii*; Sp, *Spirogyra pratensis*. (b) GUS-staining images of 10-day-old thalli harboring *_pro_*Mp*LYK1:GUS*, *_pro_*Mp*LYK2:GUS*, *_pro_*Mp*LYR:GUS*, and *_pro_*Mp*LYP:GUS*, respectively. The section is between the dorsal side and the ventral side containing the air chamber. (c) Plasma-membrane localization of MpLYK1-mCitrine and MpLYR-mTurquoise2. Magnified images of the boxed regions are also shown. Single confocal images of *M*. *polymorpha* thallus cells expressing MpLYK1-mCitrine or MpLYR-mTuquoise2. Green, cyan, and blue pseudo-colors indicate the fluorescence from mCitrine, mTurquoise2, and chlorophyll, respectively. Bars = 50 μm in wide images and 10 μm in magnified images. Note that the cell wall of air pore cells emitted autofluorescence, which is difficult to distinguish from the fluorescence of mTurquoise2 in our experimental condition.

To investigate the contribution of LysM receptor homologs to chitin and PGN responses in *M. polymorpha*, we established disruptant mutants by homologous recombination or CRISPR/Cas9-based genome editing (Supplementary Figure S5). Obvious developmental defects were not observed for the disruptant mutants under our standard growth conditions. Both chitin- and PGN-induced ROS burst were abolished in Mp*lyk1^ko^*and Mp*lyr^ge^* mutants but not in Mp*lyk2^ko^* and Mp*lyp^ko^* mutants (Figure 3a, 3b, and Supplementary Figure S7), which could be restored by expression of MpLYK1 and MpLYR under their own promoters in the respective mutants (Figure 3a, 3b, Supplementary Figure S6 and S7). Likewise, in the Mp*lyk1^ko^* and Mp*lyr^ge^* mutants, chitin-induced expression of defense-related genes, selected from the transcriptome data, was abolished (Figure 3c). Subcellular localization of fluorescent protein-tagged MpLYK1 and MpLYR indicated roles at the cell surface (Figure 2c). These results suggest that MpLYK1 and MpLYR function together to sense chitin and PGN and to activate intercellular signaling leading to defense-related gene expression.

**Figure 3.**
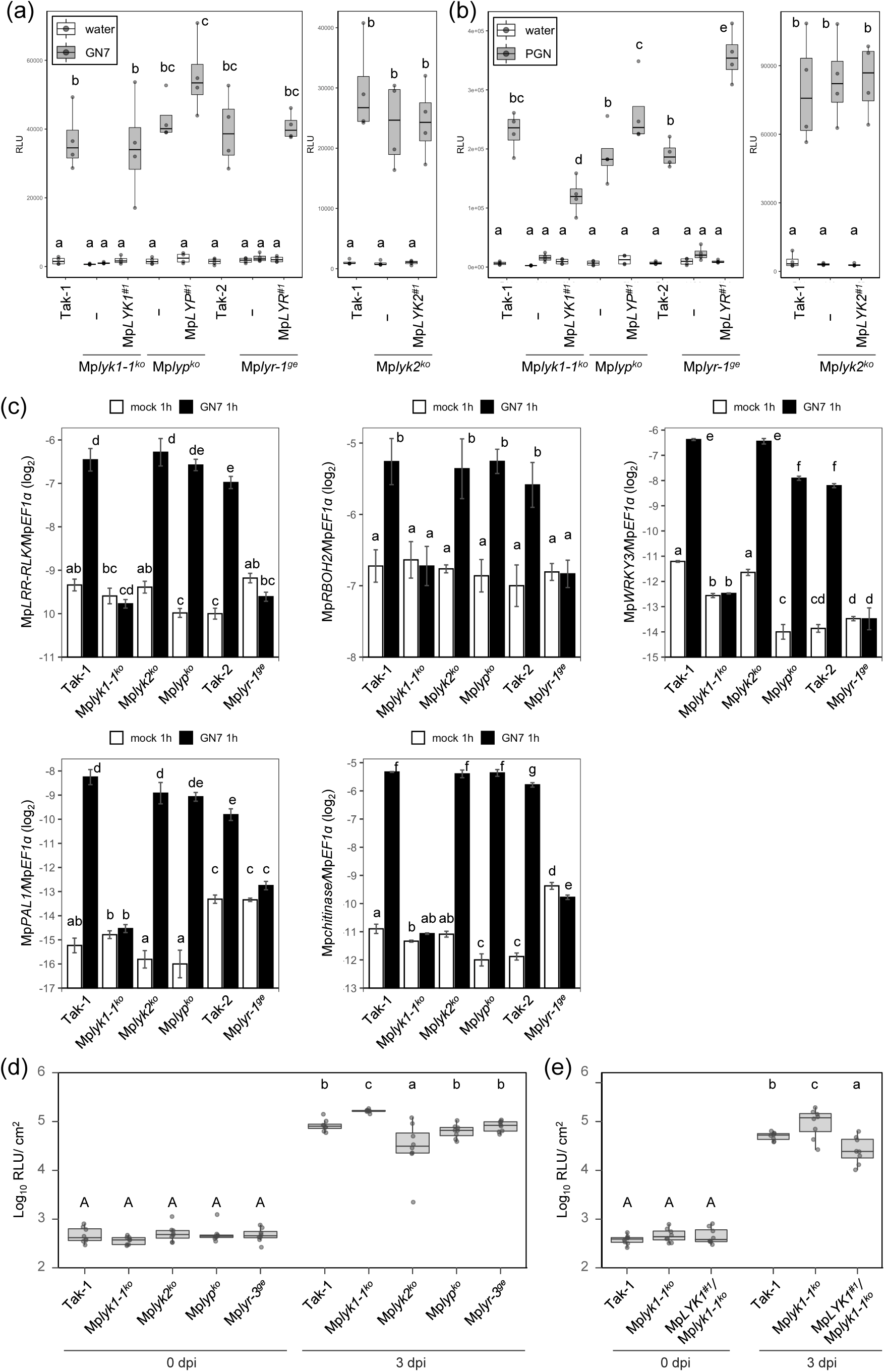
MpLYK1 and MpLYR are required for chitin- or PGN-induced responses. (a, b) Chitin or PGN-induced ROS burst in LysM receptor homolog disruptants. Six-day-old gemmalings of wild-type plants, disruptants, and complementation lines were treated with 1 μM GN7 (a) or 500 μg/ml PGN from *Bacillus subtilis* (b). The boxplot indicates total value of RLU measured by a luminometer for 30 min after GN7 treatment (a) or for 120 min after PGN treatment. Boxes show upper and lower quartiles of the value, and black lines represent the medians. Statistical groups were determined using Tukey HSD test for four replicates. Statistically significant differences are indicated by different letters (p < 0.05) (c) Chitin- induced marker gene expression in wild-type plants and disruptants. Six-day-old gemmalings were treated with 1 μM GN7 or mock for 1 hour. Mp*LRR-RLK*: Mp2g23700.1, Mp*RBOH2*: Mp3g20340.1, Mp*WRKY3*: Mp5g05560.1, Mp*PAL1*: Mp7g14880.1, Mp*chitinase*: Mp4g20440.1. Data are shown as the mean ± SE. Statistical groups were determined using the Tukey HSD test for four replicates. Statistically significant differences are indicated by different letters (p < 0.05). (d, e) Quantification of bacterial growth in the basal region of thalli, inoculated with the bioluminescent *Pto*-lux (n = 8). dpi, days post-inoculation. Statistical analysis was performed using Student t-test with p-values adjusted by the Benjamini and Hochberg (BH) method. Statistically significant differences are indicated by different letters (p < 0.05).

Air chambers have been shown to support colonization by invading microbes in *M. polymorpha* (Carella et al., 2018; Iwakawa et al., 2021). In particular, assimilatory filaments, which are specialized cell types located in air chambers for photosynthesis with less pronounced cuticle coverage, can be primarily targeted by pathogenic bacteria (Ishizaki et al., 2015). GUS reporter-based promoter analysis indicated that Mp*LYK1* and Mp*LYR* are primarily expressed in assimilatory filaments and upper epidermis (Figure 2b), consistent with their potential roles in PTI in *M. polymorpha*. Indeed, the Mp*lyk1^ko^* mutant displayed hyper-susceptibility to the pathogenic bacterium *Pseudomonas syringae* pv. *tomato* DC3000 (*Pto* DC3000) (Figure 3d and 3e). In this connection, At*cerk1* mutants are known to be hyper-susceptible towards *Pto* DC3000 (Ishizaki et al., 2013). Taken together, our results demonstrate that LysM-mediated PTI is well-conserved in the liverwort.

### Phosphoproteomic analysis of LysM-mediated signaling pathway in *M. polymorpha*

To explore downstream signaling components of LysM receptors in *M. polymorpha*, we performed differential phosphoproteomics upon chitin treatment. In total, we identified 218 proteins that were phospho-regulated 10 minutes after chitin treatment (Supplementary Table S5), a time point at which the maximum level of MAPK dual phosphorylation was observed (Figure 1d). As a proof of concept, upon chitin treatment, we observed phospho-regulation of MpLYK1 and MpLYR and the dual phosphorylation of MpMPK1, which is the only MAPK orthologous to AtMPK3, AtMPK4, and AtMPK6 (Figure 4a, 4b, 4c, and Supplementary Table S5) (Ishizaki et al., 2008). Strikingly, homologs of PTI-related components described in angiosperms are rather comprehensively identified as chitin-induced phospho-regulated proteins (Figure 4a, 4b, and Supplementary Table S5), which includes homologs of RLCKs (receptor-like cytoplasmic kinases), RBOH (respiratory burst oxidase homolog), OSCA (reduced hyperosmolality-induced [Ca^2+^] increase), MAPKKK (MAPKK kinase), MAPKK (MAPK kinase), MKP (MPK phosphatase), PP2C (protein phosphatase type 2C), WRKY, and CAMTA (calmodulin-binding transcription activator). Secretion- and autophagy-related components were also identified. This finding suggests that the intracellular signaling mechanisms leading to defense responses are also conserved in the liverwort.

**Figure 4.**
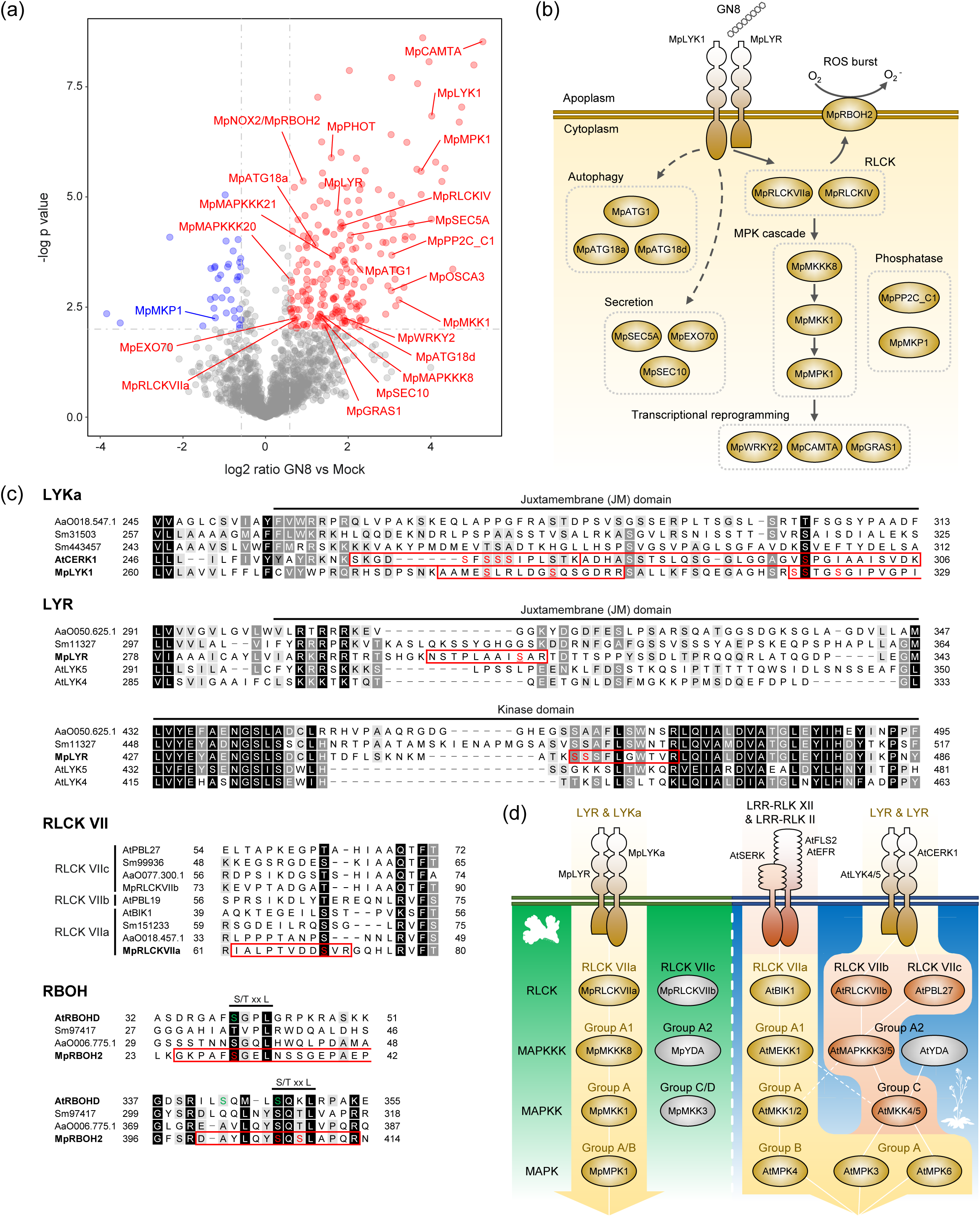
Chitin-induced phosphoproteome changes in *M. polymorpha*. (a) Volcano plots showing differential abundance of phosphopeptides between *M. polymorpha* gemmalings treated with 1 μg/ml GN8 or mock. Each dot represents a single unique phosphopeptide. Significantly increased and decreased phosphopeptides are colored red and blue, respectively (|Log_2_FC| > 0.58, p < 0.01). (b) Predicted chitin signaling pathway in *M. polymorpha* based on the identified phospho-regulated proteins. (c) Conservation and diversification of the identified phospho-sites. Identified phosphopeptides are marked with red box. Predicted phospho-sites are colored red. Confidently localized phospho-sites are further underlined. Phospho-site information on AtCERK1 is based on (Suzuki et al., 2019). Serine residues colored green in AtRBOHD are targets of AtBIK1 reported in (Kadota et al., 2014). Abbreviations: At, *Arabidopsis thaliana*; Sm, *Selaginella moellendorffii*; AaO, *Anthoceros agrestis* [Oxford]; Mp, *Marchantia polymorpha*. (d) Schematic representation of PRR signaling pathways in *M. polymorpha* and *A. thaliana*.

### Phototropin is required for repressing the induced defense-related genes in *M. polymorpha*

Our phosphoproteome profiling identified various components for which roles in PTI have not yet been described, including the blue-light receptor phototropin MpPHOT (Figure 4a and Supplementary Table S5). Phototropins from various plant species including MpPHOT are known to be activated by blue-light irradiation through induction of auto-phosphorylation, which can be visualized by a phosphorylation-dependent mobility shift on SDS-PAGE (Sugano et al., 2018). Blue-light irradiation but not chitin treatment induced a mobility shift of MpPHOT (Figure 5a), indicating that these two stimuli induce phosphorylation at different sites on MpPHOT. Indeed, differential phosphoproteomics upon blue-light irradiation revealed that different sites are phosphorylated upon chitin treatment and upon blue-light irradiation (Figure 5b and Supplementary Table S6). To investigate a potential role of MpPHOT in PTI, we compared the chitin-induced transcriptional response of the Mp*phot^ko^* mutant (female) and wild-type Tak-2 (female). K-mean clustering of DEGs identified a gene group, Cluster 6, that is uniquely up-regulated in the Mp*phot^ko^*mutant 24 hours after chitin treatment (Figure 5c). GO analysis of genes in cluster 6 revealed that defense-related genes are upregulated in the Mp*phot^ko^* mutant (Figure 5d). Expression kinetics of the cluster 6 genes indicated that MpPHOT is required for switching off gene expression during recovery from immune activation (Figure 5e). Besides, chitin-induced ROS burst was slightly upregulated in the Mp*phot^ko^* mutant (Figure 5f). Correspondingly, the Mp*phot^ko^* mutant displayed enhanced resistance to *Pto* DC3000 (Figure 5g). These results suggest a potential role of phototropin in optimal recovery of plants from unwanted long-term immune activation. The *M. polymorpha* system established here and the reported transcriptome and phosphoproteome datasets could contribute to further dissection of molecular mechanisms of PTI in plants.

**Figure 5.**
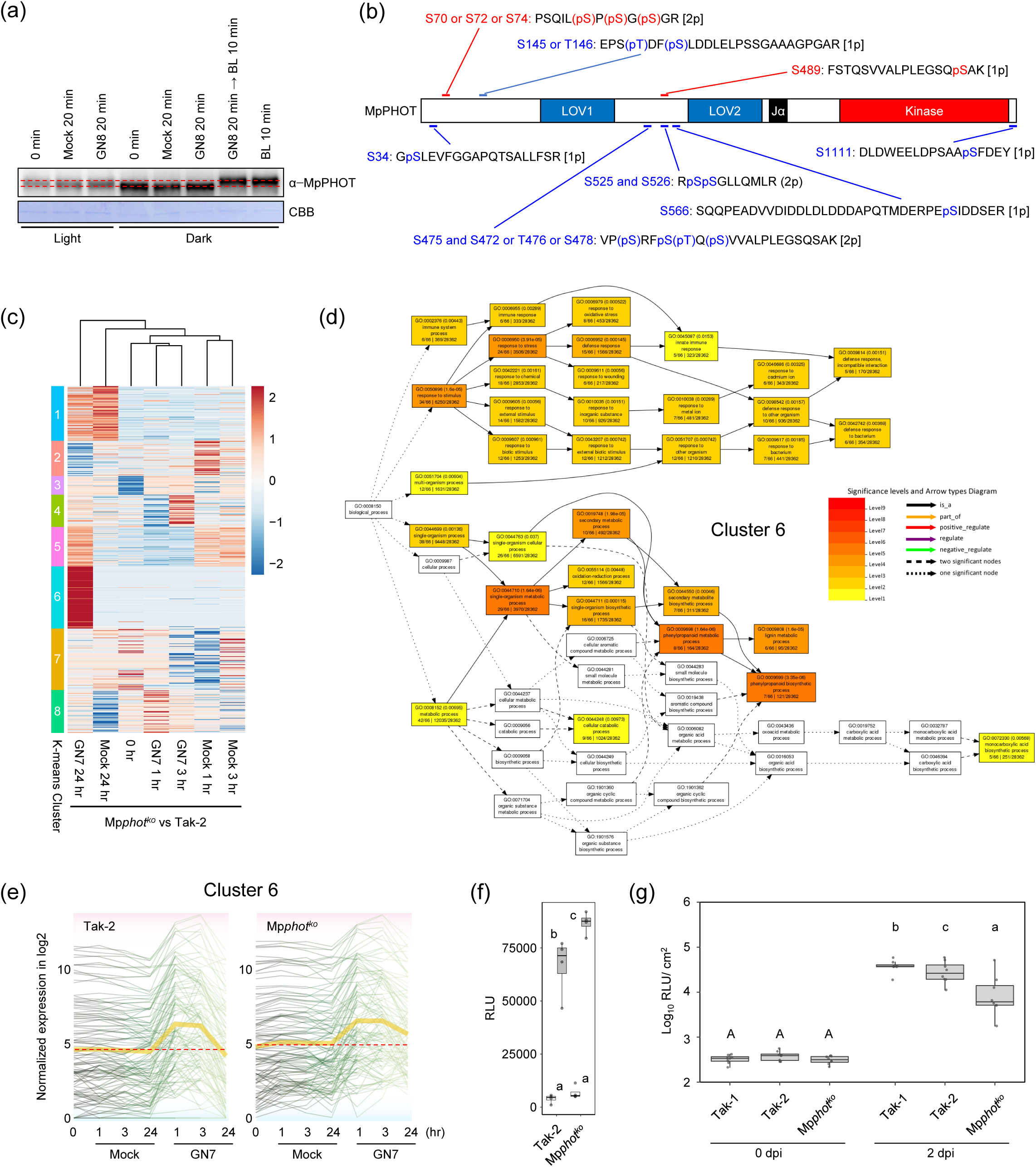
MpPHOT plays a role in PTI in *M. polymorpha*. (a) Western blot analysis of MpPHOT upon 1 μg/ml GN8 treatment or blue-light (BL) irradiation under the light or dark condition. (b) Phosphopeptides from MpPHOT induced upon GN8 treatment or blue-light irradiation identified by phosphoproteomics. Phosphopeptides induced by GN8 treatment and blue-light irradiation are colored red and blue, respectively. See Supplementary Table S5 and S6. (c) Clusters of *M. polymorpha* DEGs. Significantly differentially expressed genes showing over ±1 log2 fold change were grouped based on K-means clustering. Cluster IDs are shown on the left bar. The read count of genes was normalized into the range of ±2. See Supplementary Table S11. (d) Enriched GO terms in the K-means cluster 6 (Fig. 5c). (e) The transcription dynamics of the genes from the K-means cluster 6 (Fig. 5c) showing higher expression trends in GN7-treated condition. Yellow lines indicate mean values. (f) Chitin-induced ROS burst in Mp*phot^ko^*. Six-day-old gemmalings were treated with 1 μM N-acetylchitoheptaose (GN7). The boxplot indicates total value of RLU measured by luminometer for 30 min after GN7 treatment. Boxes show upper and lower quartiles of the value, and black lines represent the medians. Statistical groups were determined using Tukey HSD test for four replicates. Statistically significant differences are indicated by different letters (p < 0.05). (g) Quantification of bacterial growth in the basal region of thallus, inoculated with the bioluminescent *Pto*-lux (n = 8). dpi, days post-inoculation. Statistical analysis was performed using Student t-test with p-values adjusted by the Benjamini and Hochberg (BH) method. Statistically significant differences are indicated by different letters (p < 0.05).

## Discussion

PTI plays a vital role in angiosperms, but its significance in bryophytes remains elusive. In the moss *P. patens*, chitin treatment induces MAPK activation, defense-related gene expression, and cell wall modification in a Pp*CERK1*-dependent manner (Sugano et al., 2018). PpCERK is a LYKa-type LysM receptor homolog, which presumably functions as a chitin and PGN receptor or as a co-receptor for signal transduction. However, a contribution of Pp*CERK1* to resistance against pathogenic microbes has not yet been demonstrated. Growth of the liverwort *M. polymorpha* is inhibited by crude extracts from the bacterial and fungal pathogens, *Pto* DC3000, *Plectosphaerella cucumerina*, and *Fusarium oxysporum* (Ishizaki et al., 2013). Crude extracts and chitohexaose treatment can also induce defense-related gene expression in *M. polymorpha* (Redkar et al., 2022). Note that the analyzed defense-related genes have not been confirmed to be PTI-specific marker genes, and, significantly, genetic evidence for the existence of PRRs that sense potential MAMPs is missing in *M. polymorpha*. ROS burst plays an important role during PTI in angiosperms, which can be a good readout for investigating PTI-related components. However, MAMP-induced ROS burst in bryophytes has not yet been reported. Our establishment of a robust ROS burst monitoring method, which utilizes clonal gemmae, and the identification of chitin and PGN fragments as ROS burst-triggering MAMPs will be instrumental in unraveling PTI pathways in the liverwort model *M. polymorpha*. Indeed, using this system we were able to identify MpLYK1 and MpLYR as potential PRRs that are required for sensing chitin and PGN in *M. polymorpha* (Figure 3a and Supplementary Figure S8). In order to define whether MpLYK1 and/or MpLYR function as bona fide MAMP receptors, further biochemical study is required. By characterizing the chitin-induced transcriptional response and through the use of Mp*lyk1^ko^* and Mp*lyr^ge^* mutants, we were able to identify PTI-specific marker genes. Our finding that Mp*LYK1* is required for resistance against infection by the bacterial pathogen *Pto* DC3000 could be the first evidence demonstrating the significance of PTI in bryophytes (Figure 3d and 3e). As the Mp*lyr^ge^* mutant did not display hyper-susceptibility to *Pto* DC3000 (Figure 3d), there might be other PRRs that can detect bacterial MAMPs and function together with MpLYK1.

Well-studied PRRs such as *A. thaliana* FLS2 and EFR, which recognize flg22 and elf18, respectively, are LRR-RLKs from subfamily XII (LRR-RLK-XIIs). Genome analysis of *M. polymorpha* and a recent study on the expansion of immune receptor gene repertoires in plants revealed that LRR-RLK-XII genes have undergone expansion within each plant species, but with no apparent FLS2 and EFR homologs in many species including *M. polymorpha* (Ngou et al., 2022). Therefore, it is not surprising that flg22 and elf18 did not induce ROS burst in *M. polymorpha* (Figure 1a). It would be interesting to investigate whether LRR-RLK-XIIs in *M. polymorpha* also function as PRRs by sensing unidentified MAMPs.

In contrast to LRR-RLK-XIIs, LysM receptors are found to be rather well-conserved across entire land plant lineages. Our phylogenetic analysis based on ectodomain of LysM receptor homologs indicated that LysM receptors of land plants can be classified into four subgroups (Figure 2a and Supplementary Figure S4). The liverwort *M. polymorpha* and the hornworts, *Anthoceros agrestis* and *Anthoceros punctatus*, have a single gene in each subgroup. LysM receptor homologs of Charophyceae, *Chara braunii* and *Nitella mirabilis*, form an independent subgroup (Figure 2a and Supplementary Figure S4). This implies that the last common ancestor of land plants was equipped with a LysM receptor, which might have originated from a streptophyte algae LysM receptor, and suggest that the ancestral LysM receptor was duplicated and sub-functionalized early after terrestrialization and before the emergence of the diversified land plant lineages. Based on studies in angiosperms and considering that LYR-type receptors presumably lack kinase activity, LYRs, including MpLYR, may generally contribute to ligand perception. Concomitantly, LYKa may generally function as a co-receptor for intracellular signal transduction. In *A. thaliana* and rice, several LYPs function as PGN receptors (Liu et al., 2012; Willmann et al., 2011), whereas, in *M. polymorpha*, we found that MpLYP is not required for PGN-induced ROS burst (Figure 3a and Supplementary Figure S8). Along the same lines, *P. patens* has lost *LYP* but is able to sense PGN. In different plant lineages, LYPs have sub-functionalized to gain or lose PGN-binding ability. Other LYPs, i.e., OsCEBiP and AtLYM2, function as chitin receptors (Faulkner et al., 2013; Kaku et al., 2006). AtLYM2 does not play a role in conventional PTI responses but regulates chitin-induced plasmodesmata (PD) closure (Faulkner et al., 2013). Expression of Mp*LYP* in storage cells may suggest that MpLYP plays a distinct role compared to MpLYKs and MpLYR. It would be interesting to investigate whether MpLYP plays a role in PD regulation as AtLYM2. LYKb-type receptors seem to have undergone less expansion but have been retained in most plant species with the exception of mosses. Except for a few studies that have described roles of AtLYK3 in the crosstalk between immunity and responses induced by Nod factors or abscisic acid, the molecular functions of LYKb are less understood (Liang et al., 2013; Paparella et al., 2014). Further characterization of MpLYK2 may help to uncover the fundamental role of LYKb in plants.

LysM receptors also play a key role in symbiosis establishment. It is thought that the last common ancestor of land plants could establish mutualism with arbuscular mycorrhizal (AM) fungi (Rich et al., 2021). Although *M. polymorpha* is a non-mycorrhizal plant, liverworts in Marchantiales including *Marchantia paleacea* can accommodate AM fungi (Humphreys et al., 2010). Smooth rhizoids of liverworts were shown to be an entry point of AM fungi (Russell and Bulman, 2005). We found that Mp*LYK* genes are also expressed in the smooth rhizoids and rhizoid precursor cells (Supplementary Figure S8). This observation suggests a potential role of LysM receptors in AM symbiosis in Marchantiales. Analysis of LysM receptor homologs in *Marchantia paleacea* would resolve this possibility.

Our phosphoproteome analysis identified several proteins that putatively function downstream of MpLYK1 (Figure 4). We found that the juxtamembrane (JM) domains of MpLYK1 and MpLYR are phospho-regulated upon chitin treatment. The JM domain of AtCERK1 plays a significant role in chitin signal transduction and was shown to be phosphorylated, although the JM domain is generally less conserved at the amino acid sequence level (Suzuki et al., 2019; Zhou et al., 2020). The phospho-regulation of LysM receptor JM domain could be widely conserved regardless of the low sequence conservation. The RBOH is responsible for ROS burst during PTI in plants. In *A. thaliana*, flg22 and elf18 treatment activates AtBIK1, which is a subfamily VIIa RLCK (Supplementary Figure S10), to phosphorylate the N-terminal region of AtRBOHD, whose phosphorylation is required for MAMP-induced ROS burst (Kadota et al., 2014). AtBIK1 preferentially phosphorylates the [S/T]xxL motif (Kadota et al., 2014). We identified two serine residues in the N-terminal region of MpRBOH2 that were phosphorylated in the chitin-treated condition (Figure 4c). We found that these two phospho-sites correspond to the phospho-sites in AtRBOHD targeted by AtBIK1. These results suggest that MpRBOH2 is responsible for chitin-induced ROS burst and that the AtBIK1 homolog functions downstream of MpLYK1 to activate MpRBOH2. Consistently, we found that the only subfamily VIIa RLCK in *M. polymorpha*, MpRLCKVIIa, to be phospho-regulated upon chitin treatment. In *A. thaliana*, RLCKs that belong to the subfamily VIIb and VIIc, but not VIIa, function downstream of AtCERK1 (Bi et al., 2018; Yamada et al., 2016). However, phospho-regulation of the two remaining RLCK VIIs in *M. polymorpha* was not detected in our analysis. These results suggest that MpRLCKVIIa is responsible for MpLYK1-dependent signaling, although further genetic analysis is needed to confirm this idea. We find it intriguing that RLCK VIIa, group A1 MAPKKK, and group A MAPKK, which function downstream of LRR-RLK-type PRRs *in A. thaliana*, presumably function downstream of LysM-type PRRs in *M. polymorpha* (Figure 4d). Given that the subfamily XII LRR-RLK receptors, which may not function as PRRs in bryophytes, are less conserved in plants, it is possible that LRR-RLK-type PRRs were a tracheophyte innovation and utilized existing PTI signaling components. Considerable expansion of PRR repertoires and downstream kinases and the establishment of complex body plans may have led LysM-type PRRs to utilize other signaling components. It is also possible that *M. polymorpha* has lost signaling components and has uniquely evolved to have a very simple PTI pathway.

Phototropins function as blue-light receptors in tracheophytes as well as in bryophytes and regulate a wide range of blue-light responses (Christie, 2007; Kasahara et al., 2004; Komatsu et al., 2014). Recently, potato phototropins, StPHOT1 and StPHOT2, were shown to promote *Phytophthora infestans* infection in *Nicotiana benthamiana* (Naqvi et al., 2022). Conversely, virus-induced silencing of *N. benthamiana* phototropin genes reduced *P. infestans* colonization in *N. benthamiana* (Naqvi et al., 2022). This suggests that phototropin negatively regulates defense against oomycete pathogens in Solanaceae. Similarly, in this study, we found that the only phototropin in *M. polymorpha*, MpPHOT, negatively regulates defense against the bacterial pathogen *Pto* DC3000 (Figure 5g). Chitin-induced transcriptional reprograming was found to be transient both in *M. polymorpha* and *A. thaliana* (Figure 1e and Supplementary Figure S2a), which is not surprising because a constitutive immune activation is thought to be costly for plants. However, little is still known about the molecular mechanisms of how plants recover from immune activation. We found that chitin-induced early responses were not markedly affected in Mp*phot^ko^* (Figure 5e and 5f), indicating that Mp*phot^ko^* is not an autoimmune mutant exhibiting enhanced PTI responses. Instead, we revealed that Mp*phot^ko^* has a defect in properly switching off defense-related gene expression or eliminating induced defense-related genes at a later time points (Figure 5e). Further study is needed to unravel the molecular mechanisms underlying this intriguing phenomenon and the significance of chitin-induced phosphorylation of MpPHOT. It is also important to investigate whether phototropin-dependent regulation of defense gene expression is generally conserved in other plant species.

In summary, this study demonstrates that LysM-type PRR-dependent PTI is highly conserved in *M. polymorpha* and that *M. polymorpha* is an attractive plant model for investigating PTI in plants. It is our hope that the methods and genetic resources reported here as well as the transcriptome and phosphoproteome data will facilitate further dissection of PTI and its evolution.

## Materials and Methods

### Plant materials and growth condition

Male and female accessions of *M. polymorpha*, Takaragaike-1 (Tak-1) and Takaragaike-2 (Tak-2), respectively were used as wild-type. Plants were grown on 1/2 Gamborg’s B5 medium containing 1% agar at 22 under 50–60 µmol photons m**^-^**^2^s**^-^**^1^ continuous white fluorescent light. Six-day-old gemmalings (cultured mature gemmae) in liquid 1/2 Gamborg’s B5 medium containing 0.1% sucrose with shaking at 130 rpm were used for the ROS assay, MAP kinase assay, RT-PCR, and quantitative RT-PCR (qRT-PCR).

### ROS assay

Four 6-day-old gemmalings were incubated in water containing 100 μM 8-amino-5-chloro-7-phenylpyrido [3,4-d] pyridazine-1,4-(2H,3H) (L-012) (Wako, Japan) for 2 hours at 22 under darkness, followed by transfer to water containing different elicitors. ROS production was determined by counting photons derived from L-012-mediated chemiluminescence using NightSHADE LB985 (Berthold Technologies, Germany) or SpectraMax i3 (Molecular Devices, USA).

### MAP kinase assay

Four 6-day-old gemmalings were treated with 1 μg/ml N-acetylchitooctaose (GN8) or mock, then proteins were extracted using extraction buffer (50 mM Tris-HCl (pH 7.5), 10 mM MgCl_2_, 15 mM EGTA, 100 mM NaCl, 2 mM DTT, 1 mM sodium fluoride, 0.5 mM Na_3_VO_4_, 30 mM β- glycerophosphate, 0.1% (v/v) NP-40, and cOmplete protease inhibitor cocktail EDTA-free tablet (Roche, Germany)). Phosphorylated MAPK proteins were detected by immunoblot analysis with antiphospho-p44/42 MAPK (Erk1/2)(Thr202/Tyr204) (D13.14.4E) rabbit mAb (Cell Signaling Technology, USA). The blotted membrane was stained with Coomassie Brilliant blue (CBB) to verify equal loading.

### RNA-seq analysis

Twenty to thirty 9-day-old Tak-2 and Mp*phot^ko^* plants were transferred to petri dishes with water and incubated for one day and then harvested without treatment or treated with 1 μM N-acetylchitoheptaose (GN7) for 1, 3, 24 hours or with mock for 1, 3, 24 hours. Thirty-five 8-day-old *A. thaliana* Col-8 seedlings, which were cultured in 1/2 MS liquid medium containing 0.1% sucrose at 22 under 50–60 µmol photons m**^-^**^2^s**^-^**^1^ long day condition, were transferred to petri dishes with water and incubated for one day and then harvested without treatment or treated with 100 μM N-acetylchitoheptaose (GN7) for 1, 3, 24 hours or with mock for 1, 3, 24 hours. RNA-Seq library preparation was carried out using a high-throughput RNA-Seq method (Kumar et al., 2012). The 100-bp paired-end reads were sequenced on an Illumina Hiseq 4000 platform by BGI (www.genomics.cn). The *M. polymorpha* genome files were obtained from MarpolBase (MpTak1v5.1; https://marchantia.info/download/tak1v5.1/). We combined functional annotations from JGI Phytozome (https://genome.jgi.doe.gov/portal/) and Ghost Koala KEGG (Kanehisa et al., 2016). The genome files of *A. thaliana* were obtained from JGI Phytozome. The following functional annotation sets were combined for the analyses, Carbohydrate Active Enzyme database (CAZy; (Lombard et al., 2014)), the Gene Ontology (GO; (The Gene Ontology Consortium, 2015)), Kyoto Encyclopedia of Genes and Genomes (KEGG; (Ogata et al., 1999)), and EuKaryotic Orthologous Groups (KOG; (Tatusov et al., 2003)), PFAM (Finn et al., 2016), Panther (Thomas et al., 2003), and MEROPS (Rawlings et al., 2018). MEROPS and GO terms were obtained based on KEGG, GO, PFAM, IDs using R packages KEGG.db (Carlson, 2016), GO.db (Carlson, 2019), and PFAM.db (Carlson et al., 2018). The raw reads were quality-trimmed using Fastp with default parameters (Chen et al., 2018). We performed mapping reads and counting transcripts per gene with the *A. thaliana* and *M. polymorpha* genomes using STAR (Dobin et al., 2013). The log2 fold difference of the gene expression between conditions was calculated with R package DESeq2 (Love et al., 2014). The genes with very low count were excluded (less than 10 reads summed from all conditions) for the analyses. Genes with statistical significance were selected (FDR adjusted p < 0.05). Normalized read counts of the genes were also produced with DESeq2, which were subsequently log2 transformed. Differentially expressed genes were grouped using K-means clustering with R package, pheatmap (Kolde, 2019). All procedures were orchestrated with the visual pipeline SHIN+GO (Miyauchi et al., 2016; 2017; 2018; 2020). R was used for operating the pipeline (R Core Team, 2013). The RNA-seq data used for the study are available (###### accession number here - data deposition in progress). The gene ontology (GO) enrichment analysis for DEGs was performed by Metascape (https://metascape.org) (Zhou et al., 2019) or agriGO v2.0 (http://systemsbiology.cpolar.cn/agriGOv2/) (Tian et al., 2017). For GO analysis of DEGs in *M. polymorpha*, the best BLASTP hit genes in *A. thaliana* was used (Best.hit.arabi.name_v3.1 in Supplementary Table S7 and S11).

### Database search

Amino acid sequences of LysM receptor homologs were obtained from the following databases: https://www.arabidopsis.org for *Arabidopsis thaliana*, https://lotus.au.dk for *Lotus japonicus*, https://phytozome.jgi.doe.gov/pz/portal.html#!info?alias=Org_Mtruncatula for *Medicago truncatula*, http://groups.english.kib.cas.cn/epb/dgd/Download/201711/t20171101_386248.html for *Cuscuta australis*, https://www.plabipd.de/project_cuscuta2/start.ep for *Cuscuta campestris*, https://phytozome.jgi.doe.gov/pz/portal.html#!info?alias=Org_Smoellendorffii for *Selaginella moellendorffii*, https://phytozome.jgi.doe.gov/pz/portal.html#!info?alias=Org_Sfallax for *Sphagnum fallax*, https://phytozome.jgi.doe.gov/pz/portal.html#!info?alias=Org_Ppatens for *Physcomitrium patens*, https://marchantia.info for *Marchantia polymorpha*, and https://bioinformatics.psb.ugent.be/orcae/overview/Chbra for *Chara braunii*. *LysM* genes in *Oryza sativa* were previously described in (Liu et al., 2012; Shimizu et al., 2010). The *LysM* gene in *Nitella mirabilis* was previously described in (Delaux et al., 2015).

### Phylogenetic analysis of LysM receptor homologs

Alignment of full-length proteins was constructed using MUSCLE alignment implemented in the Geneious 9.1.2 software package (Biomatters; http://www.geneious.com) at the default parameters, and then LysM1, LysM2, and LysM3 domains, which were previously defined in (Madsen et al., 2003), were extracted from the full-length protein alignment. An unrooted or rooted maximum-likelihood phylogenetic tree was constructed using PhyML program ver. 2.2.0 (Guindon and Gascuel, 2003) implemented in the Geneious software, using the LG as substitution model.

### Genomic DNA extraction

Total DNA was extracted from approximately 1 g of fresh weight of 6-day-old gemmalings using the Cetyl trimethyl ammonium bromide (CTAB) method as previously described in (Kubota et al., 2014) or using ISOPLANT II (Nippon gene).

### RNA extraction and cDNA synthesis

Total RNA was extracted from 6-day-old gemmalings using the RNeasy Plant Mini Kit (QIAGEN) or NucleoSpin RNA Plant (Macherey-Nagel). First-strand complementary DNA was synthesized from 0.5 μg total RNA using a ReverTra Ace qPCR RT Master Mix with gDNA Remover (TOYOBO).

### Plasmid constructions and transformation

The 5 kbp putative promoter fragment upstream of the translation initiation codon of each gene was cloned into pENTR4 dual-selection vector (Thermo Fisher SCIENTIFIC, USA) using an In-Fusion HD cloning kit, and then they were subcloned into binary vector pMpGWB104 (Ishizaki et al., 2015) for constructing *_pro_*Mp*LYK1:GUS*, *_pro_*Mp*LYK2:GUS*, *_pro_*Mp*LYR:GUS*, and *_pro_*Mp*LYP:GUS* using LR clonase II enzyme mix. To generate the targeting vectors for Mp*lyk1-1^ko^*, Mp*lyk1-2^ko^*, Mp*lyk2^ko^*, and Mp*lyp^ko^*, homologous arms were amplified from Tak-1 genomic DNA using KOD Plus Neo (Toyobo, Japan). The PCR-amplified fragments of the 5’ end and 3’ end were cloned into the *Pac*I site and *Asc*I site of pJHY-TMp1 (Ishizaki et al., 2013), respectively, using an In-Fusion HD cloning kit (Clontech Laboratories, USA). Targeting vectors were introduced into F1 sporelings of *M. polymorpha* derived from crosses between Tak-1 and Tak-2, as described previously (Ishizaki et al., 2008). To generate the targeting vectors for Mp*lyr-1^ge^*, Mp*lyr-2^ge^*, and Mp*lyr-3^ge^*, annealed oligos for an Mp*LYR*-targeting gRNA1 and an Mp*LYR*-targeting gRNA2 were ligated into *Bsa*I-digested pMpGE_En03 (Sugano et al., 2018) using an T4 DNA ligase (NEB, UK). Mp*lyr-1^ge^*was generated using Mp*LYR*-targeting gRNA1. Mp*lyr-2^ge^* was generated using both Mp*LYR*-targeting gRNA1 and gRNA2. Mp*lyr-3^ge^*was generated using Mp*LYR*-targeting gRNA2. Mp*LYR*-targeting gRNAs were subcloned into binary vector pMpGE010 (Sugano et al., 2018) using LR clonase II enzyme mix (Thermo Fisher SCIENTIFIC, USA). Screening for homologous recombination-mediated gene-targeted lines was performed by genomic PCR as described previously (Ishizaki et al., 2013). Screening for CRISPR/Cas9-mediated targeted mutagenesis lines was performed by genomic PCR as described previously (Sugano et al., 2014). The open reading frame (ORF) fragment of Mp*LYK1* and Mp*LYK2* was cloned into corresponding pENTR4-promoter, and then they were subcloned into binary vector pMpGWB301 (Ishizaki et al., 2015) for constructing *_pro_*Mp*LYK1:*Mp*LYK1* and *_pro_*Mp*LYK2:*Mp*LYK2*. The ORF fragment of Mp*LYP* was cloned into pENTR4. The ORF fragment of Mp*LYR* was cloned into pENTR4, then the PAM sequence of Mp*LYR* in pENTR4-Mp*LYR* was mutated, as shown in Supplementary Figure S5d, using PrimeSTAR® Mutagenesis Basal Kit (TaKaRa, Japan) so as not to be targeted by CRISPR/Cas9. pENTR4-*_pro_*Mp*LYP* and pENTR4-*_pro_*Mp*LYP* were subcloned into binary vector pMpGWB301, and then the ORF fragment of Mp*LYP* and mMp*LYR* was cloned into the corresponding pMpGWB301-promoter for constructing *_pro_*Mp*LYP:*Mp*LYP* and *_pro_*Mp*LYR:*mMp*LYR*. The resultant plasmids were introduced into corresponding knockout mutants, as described previously (Ishizaki et al., 2013). The primers used are listed in Supplementary Table S3 (No. 1−46).

### Assays for GUS activity and sectioning

Histochemical GUS assays were performed according to the method described by (Jefferson et al., 1987) with some modifications, as previously described (Ishizaki et al., 2012). For sectioning, GUS-stained samples were embedded into Technovit 7100 resin according to the manufacturer’s instructions (Heraeus Kulzer). Embedded samples were then sectioned into 10 μm-thick sections using RM2125 RTS microtome (Leica, Germany) with TC-65 tungsten blade.

### Confocal laser scanning microscopy

Five-day-old gemmalings grown on 1/2× Gamborg’s B5 medium containing 1.0% (w/v) sucrose and 1.0% (w/v) agar at 22 °C under continuous white light were used for observation. The samples were mounted in a 1/2× Gamborg’s B5 liquid medium and observed using an LSM780 confocal microscope (Carl Zeiss) equipped with an oil immersion lens (63×, numerical aperture = 1.4). The plant expressing MpLYK1-mCitrine was excited at 488 nm (Argon) and 561 nm (DPSS 561-10), and emissions between 482–659 nm were collected. The plant expressing MpLYR-mTurquoise2 was excited at 405 nm (Diode 405-30)), and emissions between 428–659 nm were collected. Spectral unmixing of the obtained images was conducted using ZEN2012 software (Carl Zeiss).

### Quantitative RT-PCR or semi quantitative RT-PCR

Quantitative RT-PCR was performed using a LightCycler 96 (Roche, Basel, Switzerland). Thunderbird SYBR qPCR Mix (Toyobo) was used for amplification. Mp*EF1*_α_ was used as an internal standard. Semi quantitative RT-PCR was performed using a thermal cycler. Mp*ACT1* was used as an internal standard. Primers used for qRT-PCR and semi qRT-PCR are listed in Supplementary Table S3 (No. 47−90).

### Bioluminescence-based bacteria quantification

Bacterial quantification in infected thalli was carried out as described before (Matsumoto et al., 2022). Briefly, *M. polymorpha* were grown on autoclaved cellophane disc on half-strength GB5 medium for two weeks. In the meantime, *Pto*-lux was cultivated in King’s B medium containing 30 μg/mL rifampicin to achieve an OD_600_ of 1.0. The saturated bacterial culture was subsequently washed and resuspended in Milli-Q water to prepare a bacterial suspension with of an OD_600_ of 1.0. Next, 2-week-old thalli were submerged in the bacterial suspension followed by vacuum for 5 min and incubation for 0 to 3 days on pre-wetted filter papers. After incubation, thallus discs (5 mm diameter) were punched from the basal region using a sterile biopsy punch (pfm medical) and transferred to a 96-well plate (VWR). Bioluminescence was measured in a FLUOstar Omega plate reader (BMG Labtech).

### Phosphoproteome analysis

Ten 10-day-old Tak-2 gemmalings, which were cultured in 1/2 B5 liquid medium containing 0.1% sucrose, were transferred to petri dishes with water and incubated in the dark at 22 for 3 days. Then gemmalings were treated with 1 μg/ml N-acetylchitooctaose (GN8) or mock for 10 min in dark condition or irradiated with blue-light (90 μmol m^-2^s^-1^) (MIL-B18, SANYO Electric, Japan) for 10 min at room temperature and then immediately frozen with liquid nitrogen. Sample preparation and liquid chromatography-tandem mass spectrometry (LC– MS/MS) analysis was performed as described previously with minor modifications (Koide et al., 2020). Raw data were processed using MaxQuant software (version 1.6.3.4, http://www.maxquant.org/) (Cox and Mann, 2008) with label-free quantification (LFQ) and iBAQ enabled (Tyanova et al., 2016). MS/MS spectra were scanned by the Andromeda search engine against a combined database containing the sequences from *M. polymorpha* (MpTak1v5.1_r1.protein.fasta, https://marchantia.info/download/tak1v5.1/), sequences of 248 common contaminant proteins, and decoy sequences. Trypsin specificity was required and a maximum of two missed cleavages allowed. Minimal peptide length was set to seven amino acids. Carbamidomethylation of cysteine residues was set as fixed, phosphorylation of serine, threonine and tyrosine, oxidation of methionine and protein N-terminal acetylation as variable modifications. The match between runs option was enabled. Peptide-spectrum-matches and proteins were retained if they were below a false discovery rate of 1% in both cases. Statistical analysis of the intensity values obtained for the phospho-modified peptides (“modificationSpecificPeptides.txt” output file) was carried out using Perseus (version 1.5.8.5, http://www.maxquant.org/). Intensities were filtered for reverse and contaminant hits and the data was filtered to retain only phospho-modified peptides. Next, intensity values were log2 transformed. After grouping samples by condition only those sites were retained for the subsequent analysis that had four valid values in one of the conditions in case of GN8 vs mock analysis and two valid values in one of the conditions in case of GN8, blue-light, mock analysis. Two-sample t-tests were performed using a permutation-based FDR of 0.05. Alternatively, the valid value-filtered data was median-normalized and missing values were imputed from a normal distribution, using the default settings in Perseus (1.8 downshift, separately for each column). The Perseus output was exported and further processed using Excel and RStudio. The MS proteomics data have been deposited in the ProteomeXchange Consortium via the PRIDE partner repository with the dataset identifiers PXD038903 and PXD038907.

## Key resources table

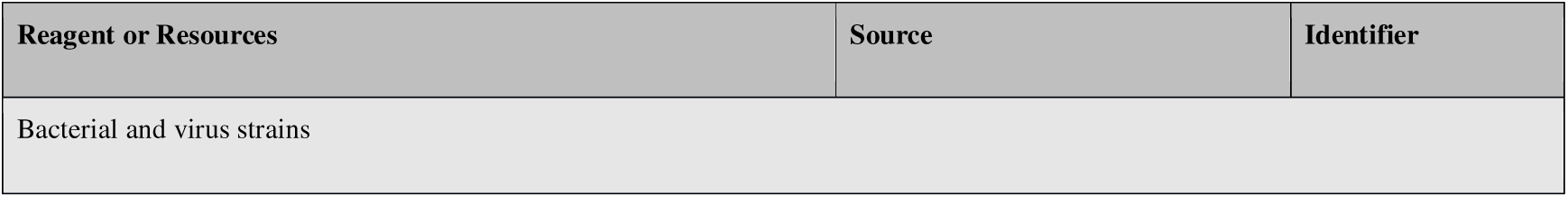
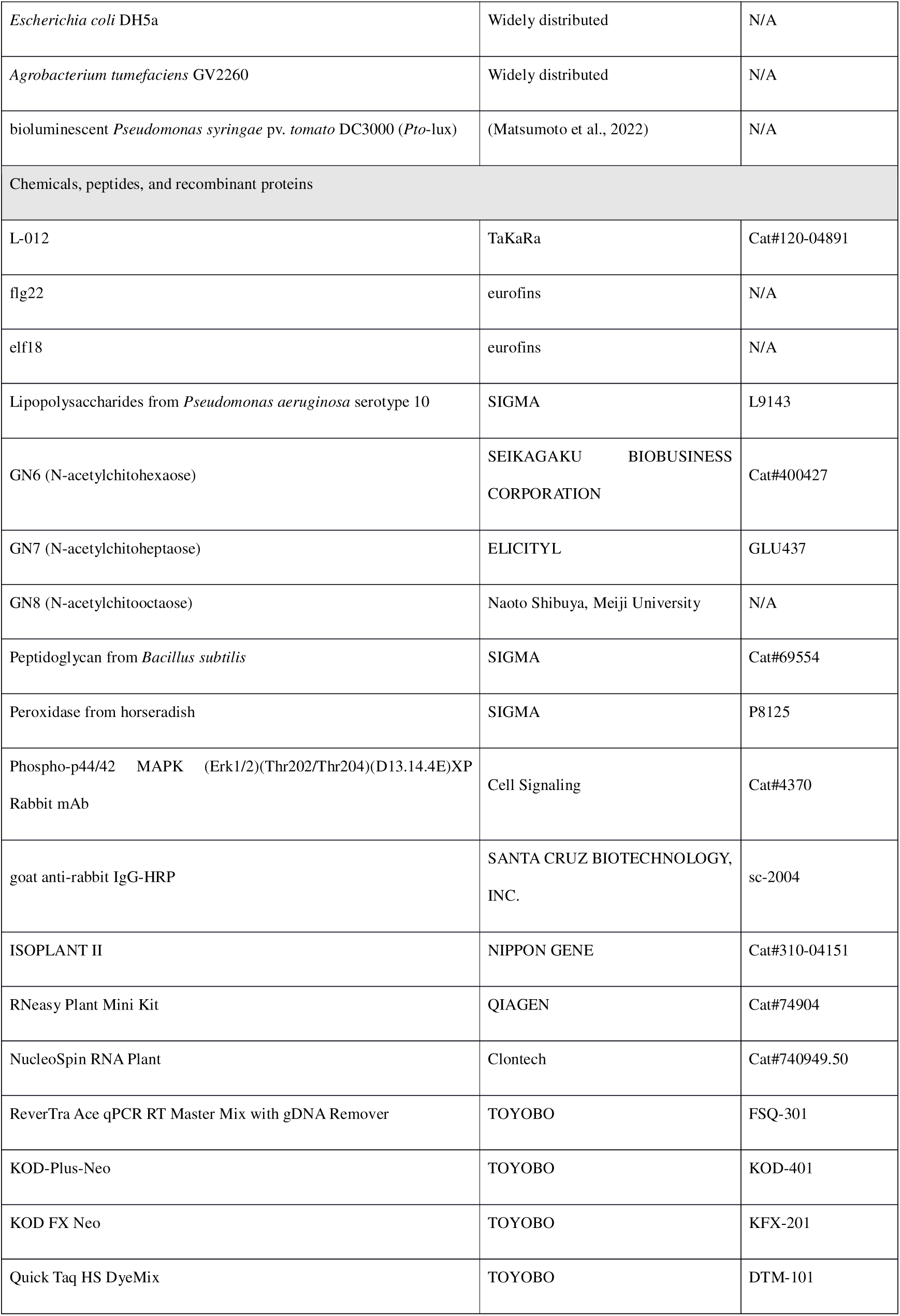
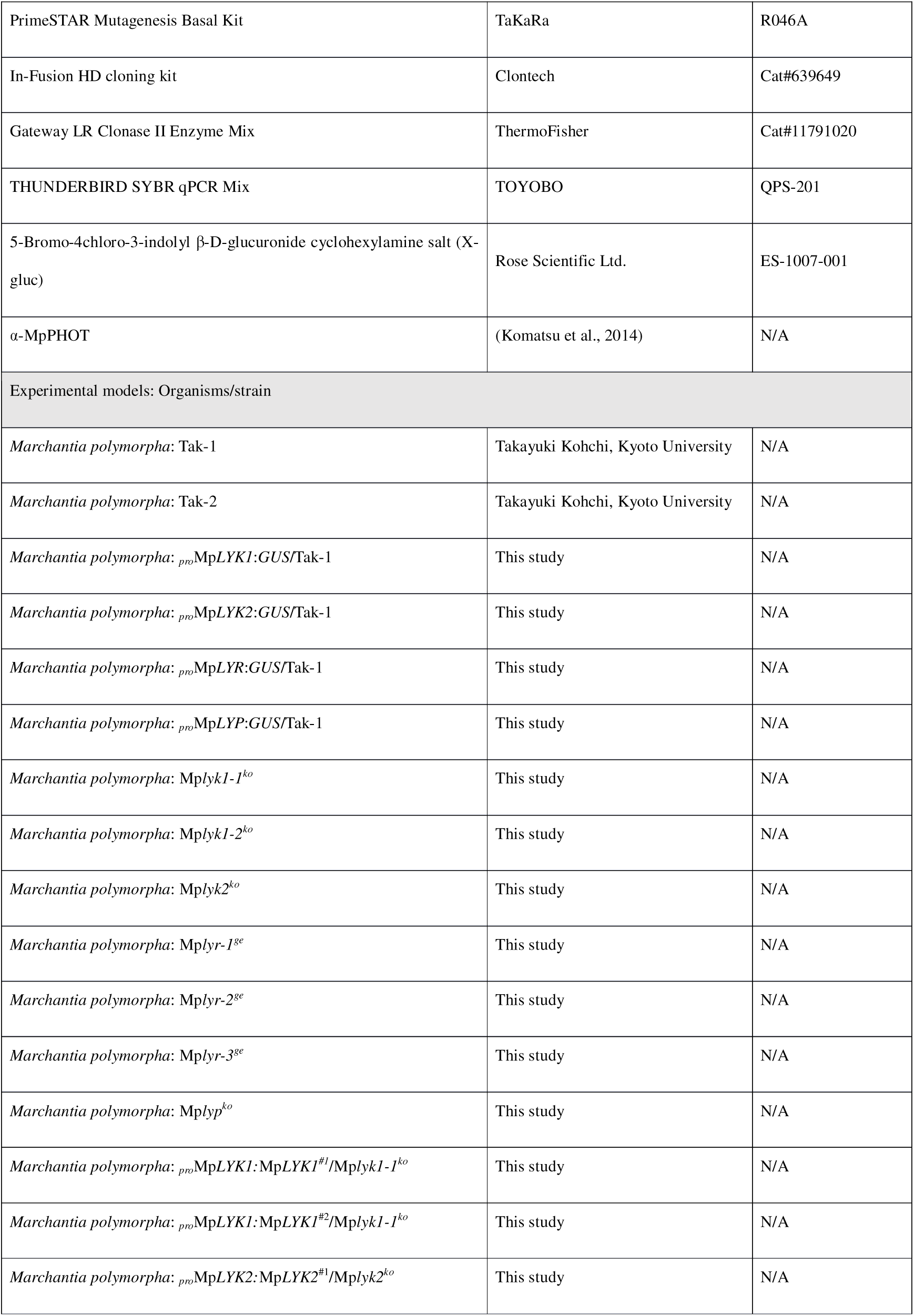
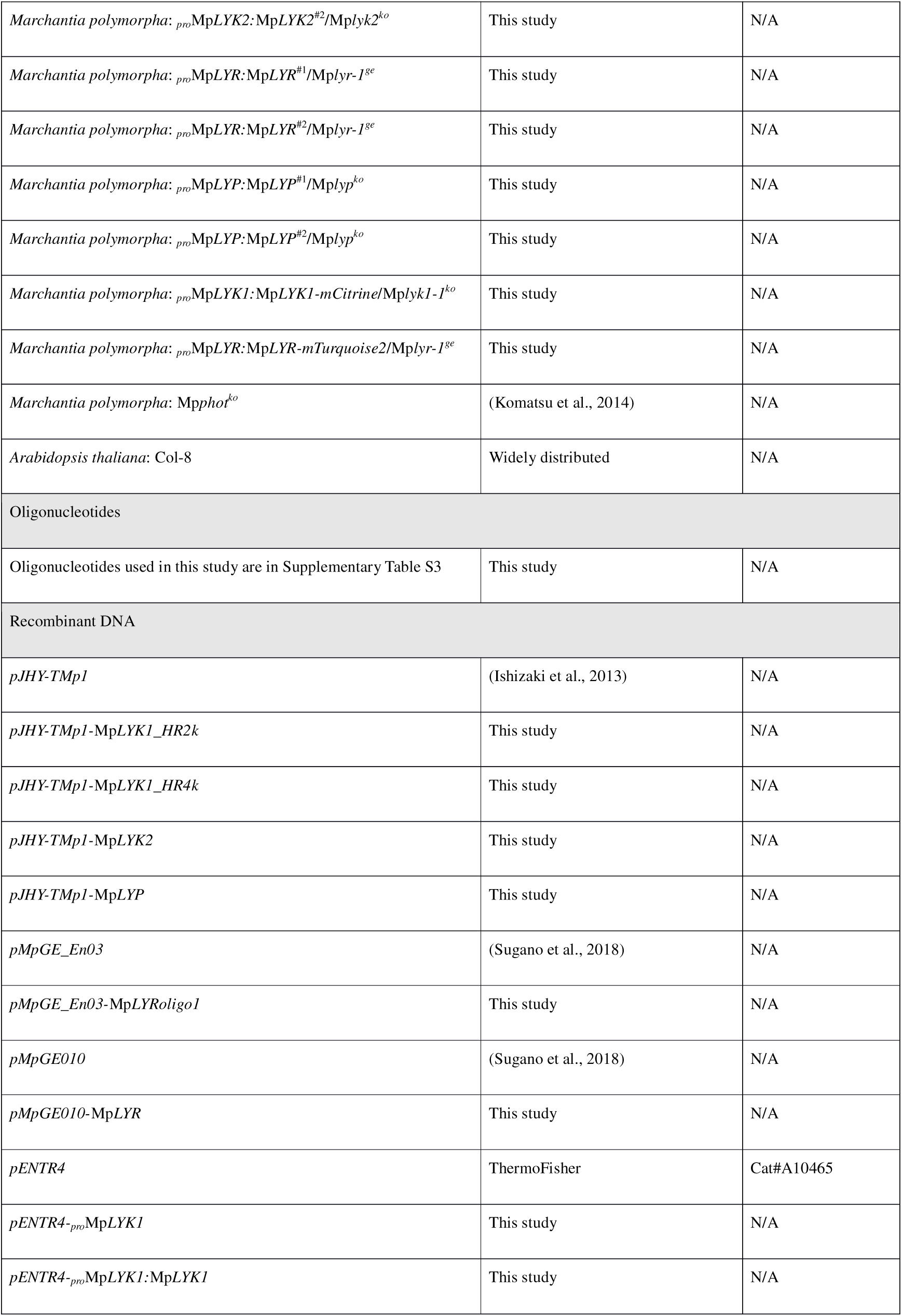
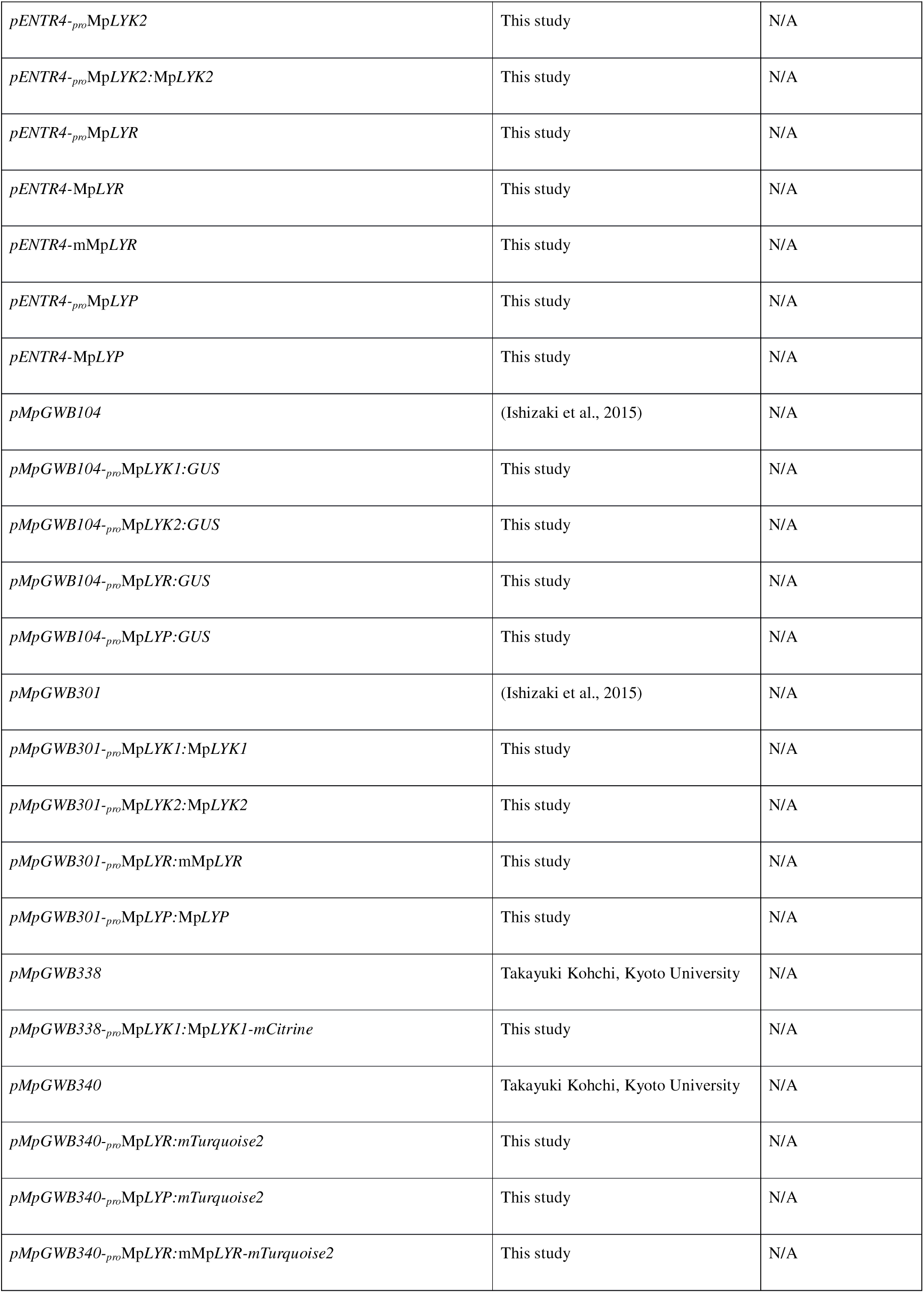

## Acknowledgments

We thank Naoto Shibuya (Meiji University, Japan) for providing N-acetylchitooctaose. We thank Neysan Donnelly (MPIPZ, Germany) for editing the manuscript.

## Author contributions

I.Y. and H.N. designed the research. I.Y., S.S., H.I., K.M., R.N., and T.Ko. generated plant materials. T.Ka., K.M., H.J., and T.U. performed microscopic analysis. T.S., S.G.I, K.M., and R.S. performed *Pto* DC3000 infection assay. I.Y., Y.I., M.S., K.S., and S.M. performed transcriptomic analysis. I.Y., Y.N., H.M., S.C.S., and H.N. performed phosphoproteomic analysis. I.Y. performed all other experiments. I.Y., H.M., and H.N. wrote the manuscript. All authors corrected the manuscript.

## Supplemental information

**Figure S1.**
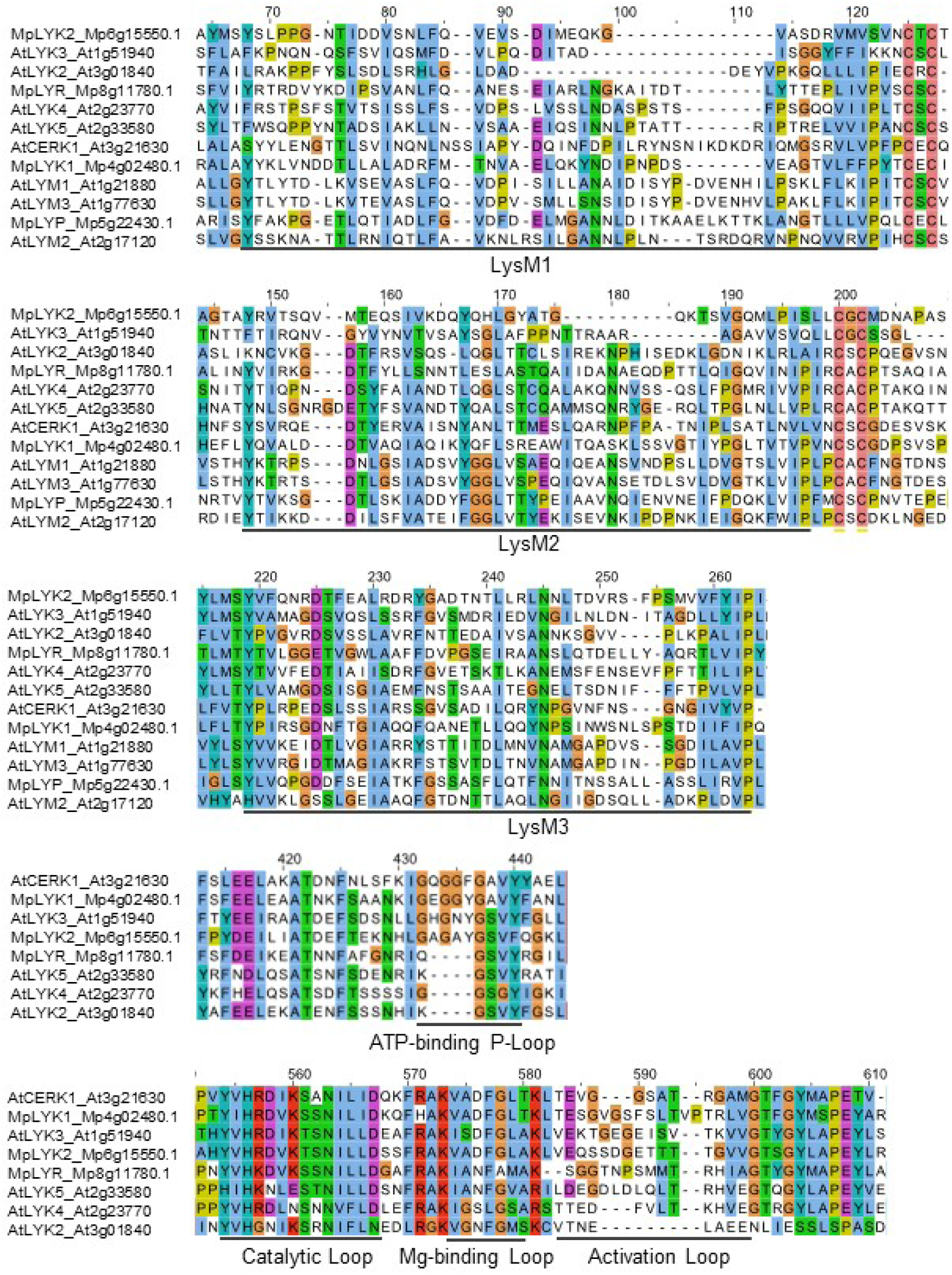
Alignment of LYKs and LYRs in *M. polymorpha* and *A. thaliana*.

**Figure S2.**
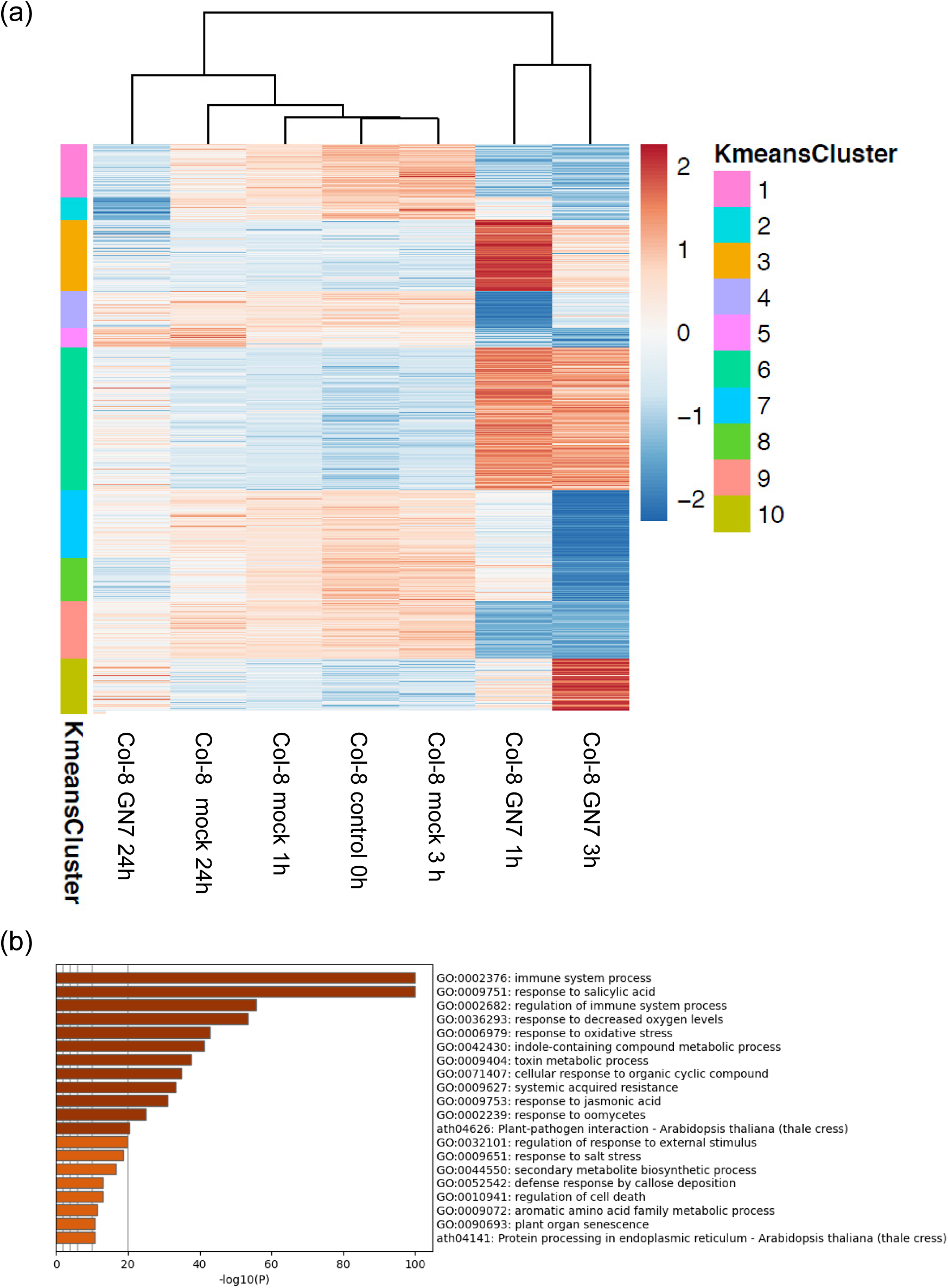
Chitin-induced transcriptional reprogramming in *A. thaliana* seedlings. (a) Clusters of *A. thaliana* DEGs. Significantly differentially expressed genes with over ±2 log2 fold changes (FDR adjusted p < 0.05) were grouped based on K-means clustering. K-means cluster ID is shown on the left bar. Log2 read count of genes was normalized into the range of ±2. See Supplementary Table S9. (b) Enriched GO terms in the *M. polymorpha* DEGs (Supplementary Figure S2a). See Supplementary Table S10.

**Figure S3.**
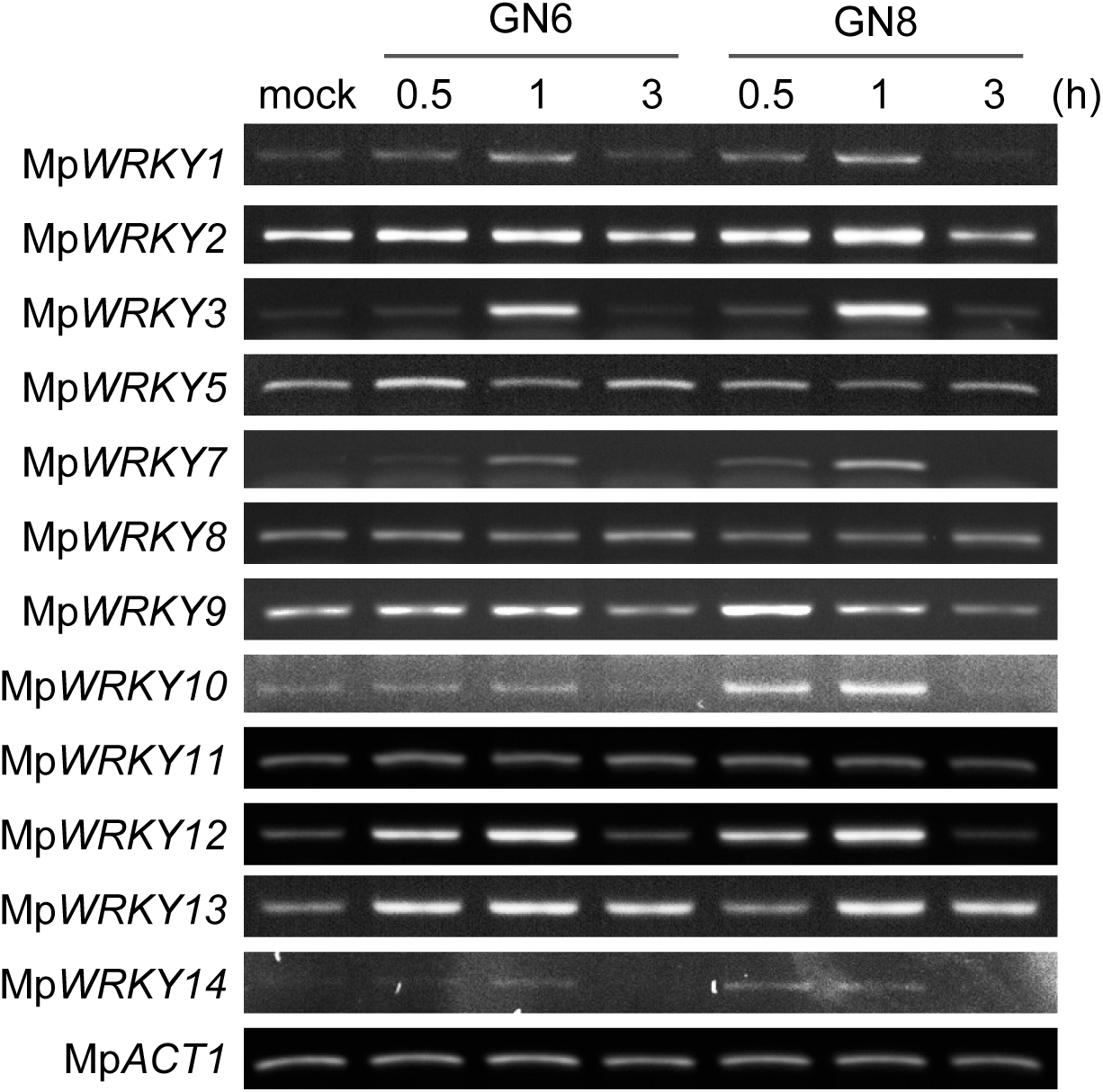
Mp*WRKY* gene expression upon chitin treatment in *M. polymorpha* Tak-1. Six-day-old gemmalings were treated with 1 μg/ml N-acetylchitohexaose (GN6) or N-acetylchitooctaose (GN8) for the indicated times. Mp*ACT1* was used as an internal control.

**Figure S4.**
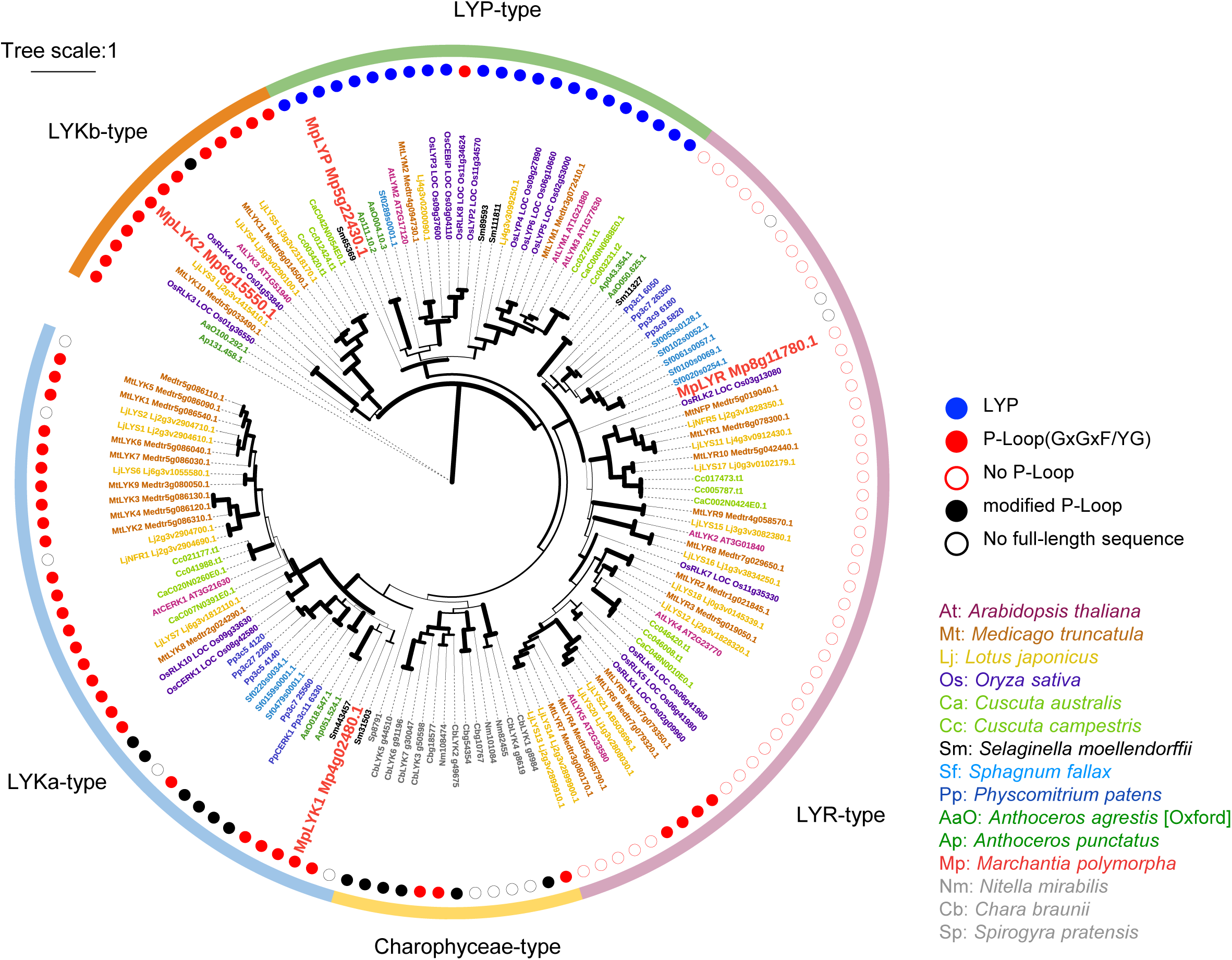
Circular phylogenetic tree of LysM proteins in plants. Circular version of the phylogenetic tree in Figure 2a.

**Figure S5.**
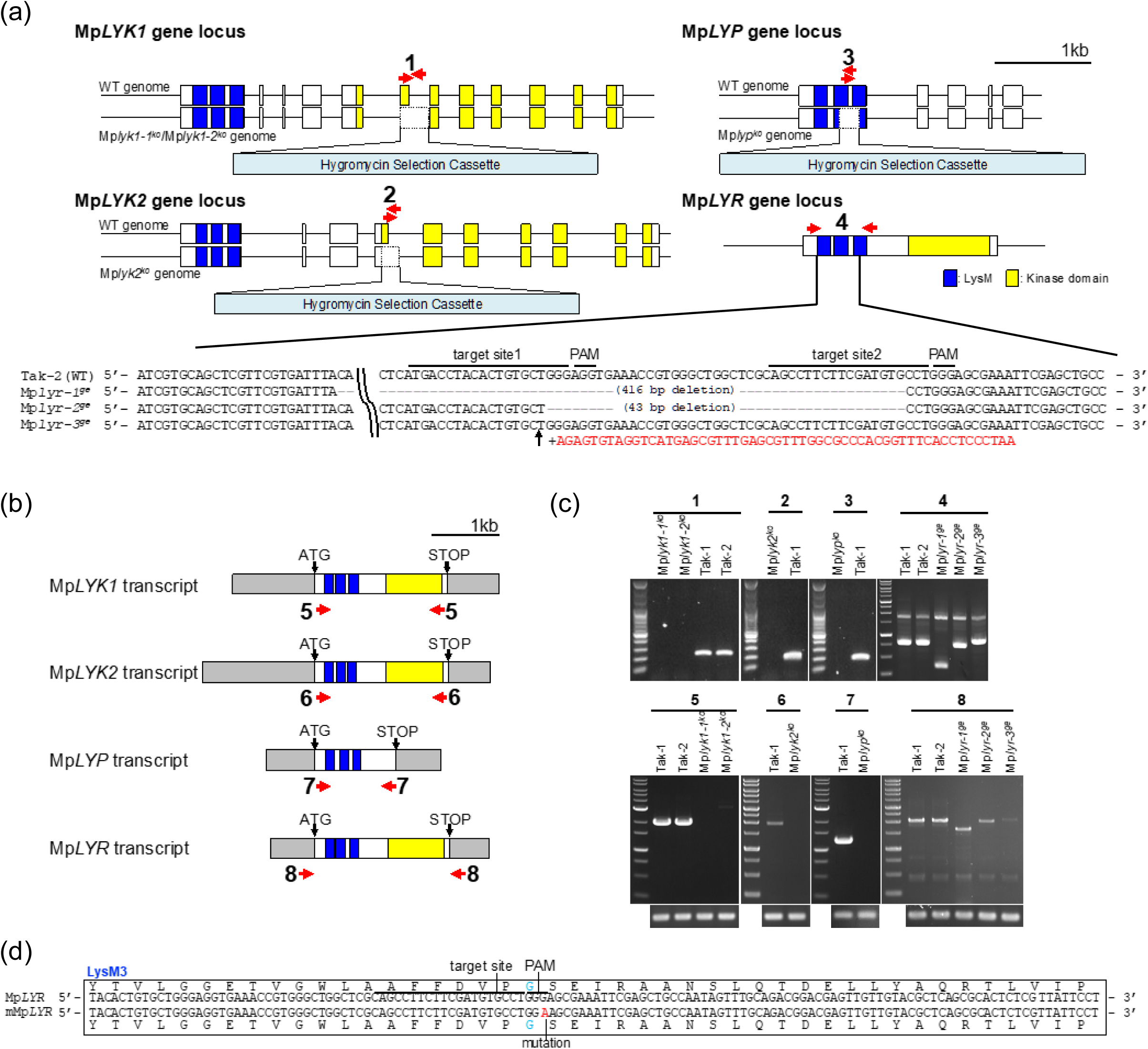
Mp*LysM* disruptants generated and used in this study. (a) Schematic representation of the targeted disruption of the Mp*LYK1*, Mp*LYK2*, and Mp*LYP* locus by homologous recombination and disruption of the Mp*LYR* locus by genome editing. All disruptants were generated using F1 sporelings of *M. polymorpha* derived from crosses between Tak-1 and Tak-2. Mp*lyk1-1^ko^*, Mp*lyk2^ko^*, Mp*lyp^ko^*, Mp*lyr-2^ge^*, and Mp*lyr-3^ge^* are male. Mp*lyk1-2^ko^* and Mp*lyr-1^ge^* are female. 416 bp or 43 bp were deleted as indicated in Mp*lyr-1^ge^* or Mp*lyr-2^ge^*, respectively. 55 bp was inserted as indicated in Mp*lyr-3^ge^*. Blue boxes and yellow boxes indicate LysM and kinase domains, respectively. Red arrows indicate primer set for amplifying the genomic region used for replacing the *hygromycin phosphotransferase* (*hptII*) cassette in Mp*LYK1*, Mp*LYK2*, and Mp*LYP* locus or the edited region in the Mp*LYR* locus. (b) The transcribed region of the *LysM* genes. Blue boxes and yellow boxes indicate LysM and kinase domains, respectively. Grey boxes indicate untranslated region. Red arrows indicate primer set for amplifying the full-length coding sequences of *LysM* genes in *M. polymorpha*. (c) The amplification of the genome region or full-length cDNA of LysM genes in WT (Tak-1and Tak-2) and disruptants using the primer sets shown in (a) and (b). (d) The mutation which did not result in an amino acid substitution so as not to be targeted by the construct directed at Mp*LYR* was inserted for Mp*LYR* complementation to Mp*lyr-1*^ge^.

**Figure S6.**
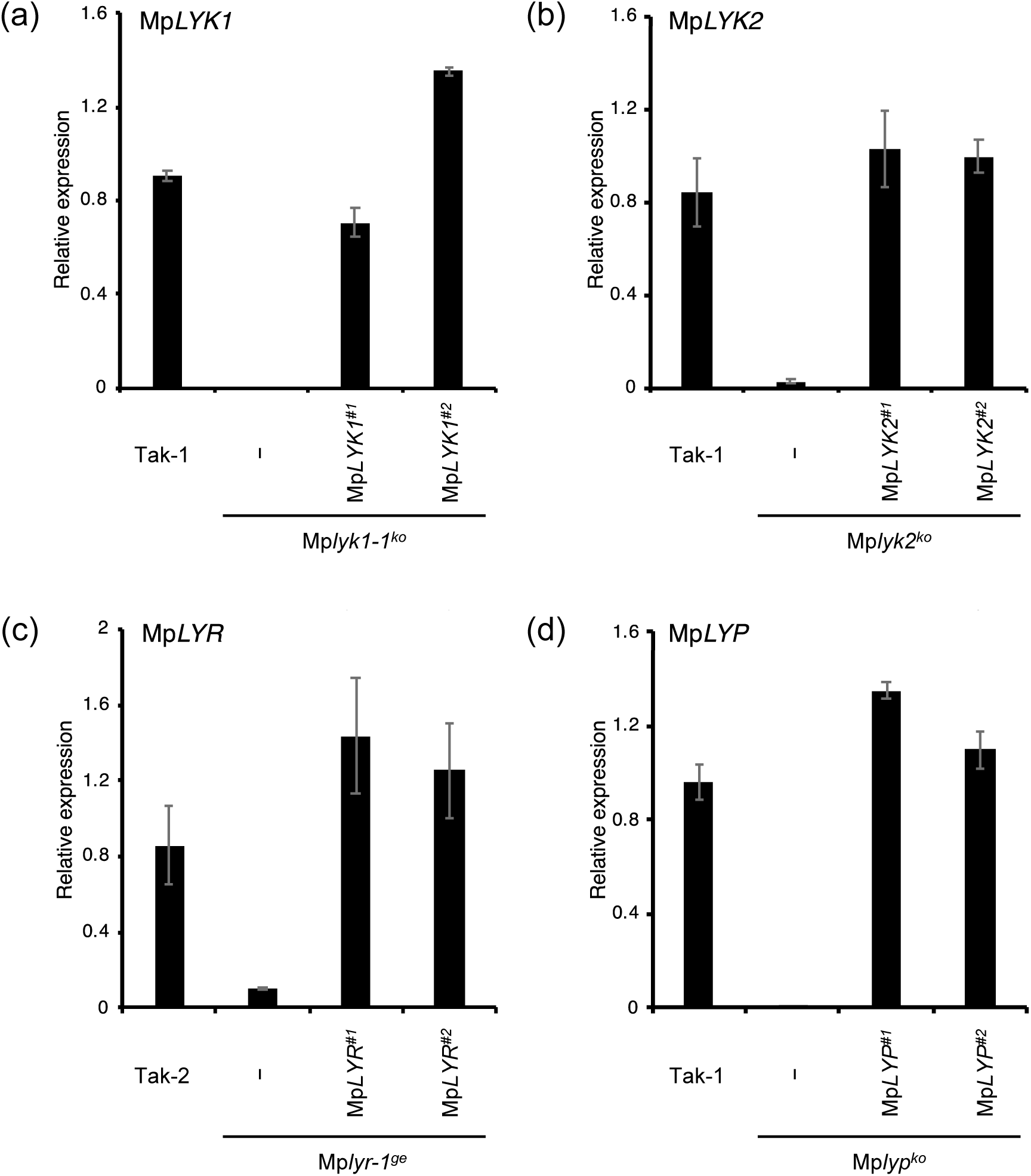
Mp*LysM* gene expression in wild-type and the transgenic plants. Mp*LYK1* (a), Mp*LYK2* (b), Mp*LYR* (c), and Mp*LYP* (d) expression in 6-day-old gemmalings grown in ½ B5 liquid medium containing 0.1% sucrose examined by qRT-PCR. The values represent the average and standard errors of three replicate experiments.

**Figure S7.**
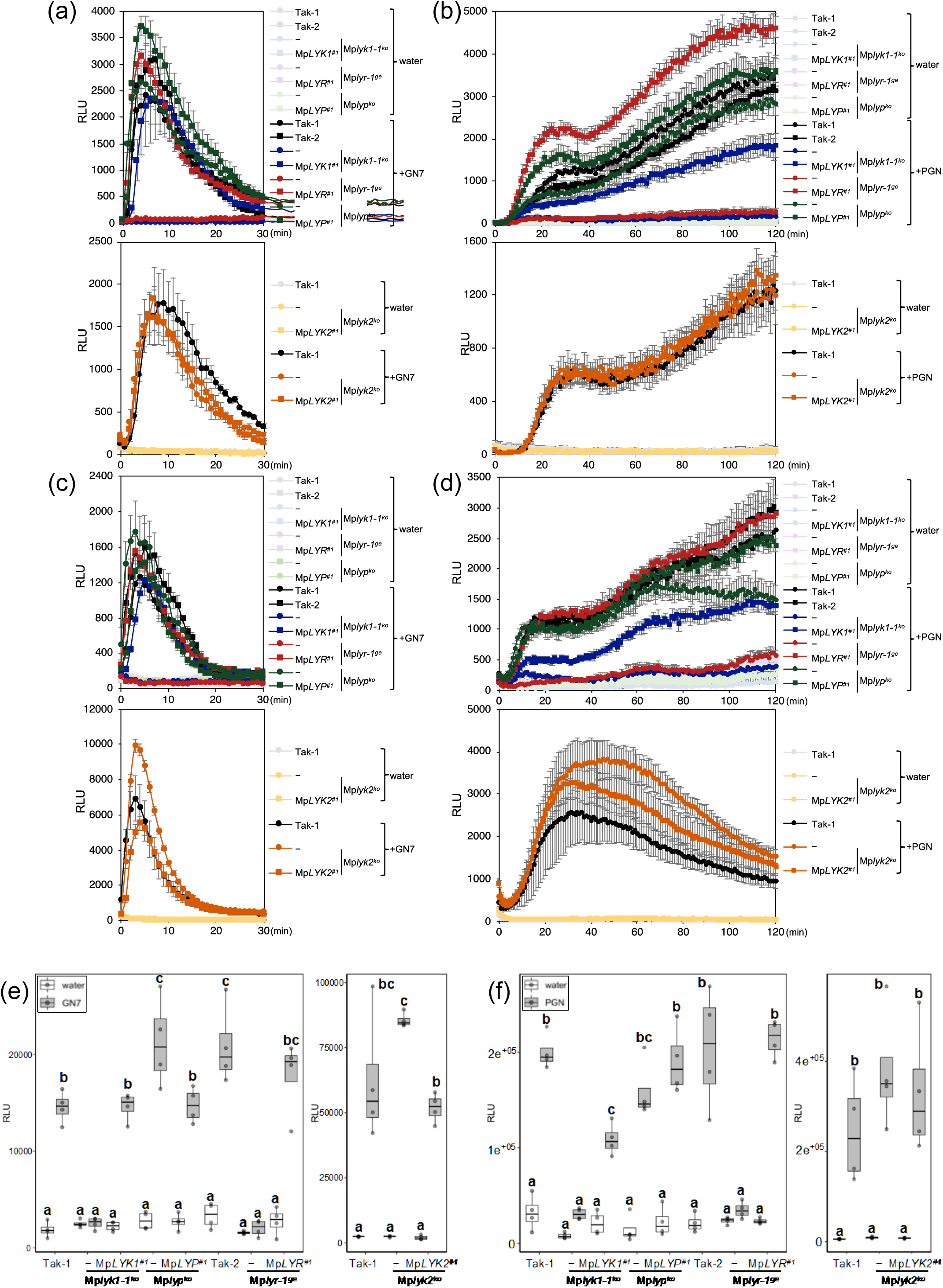
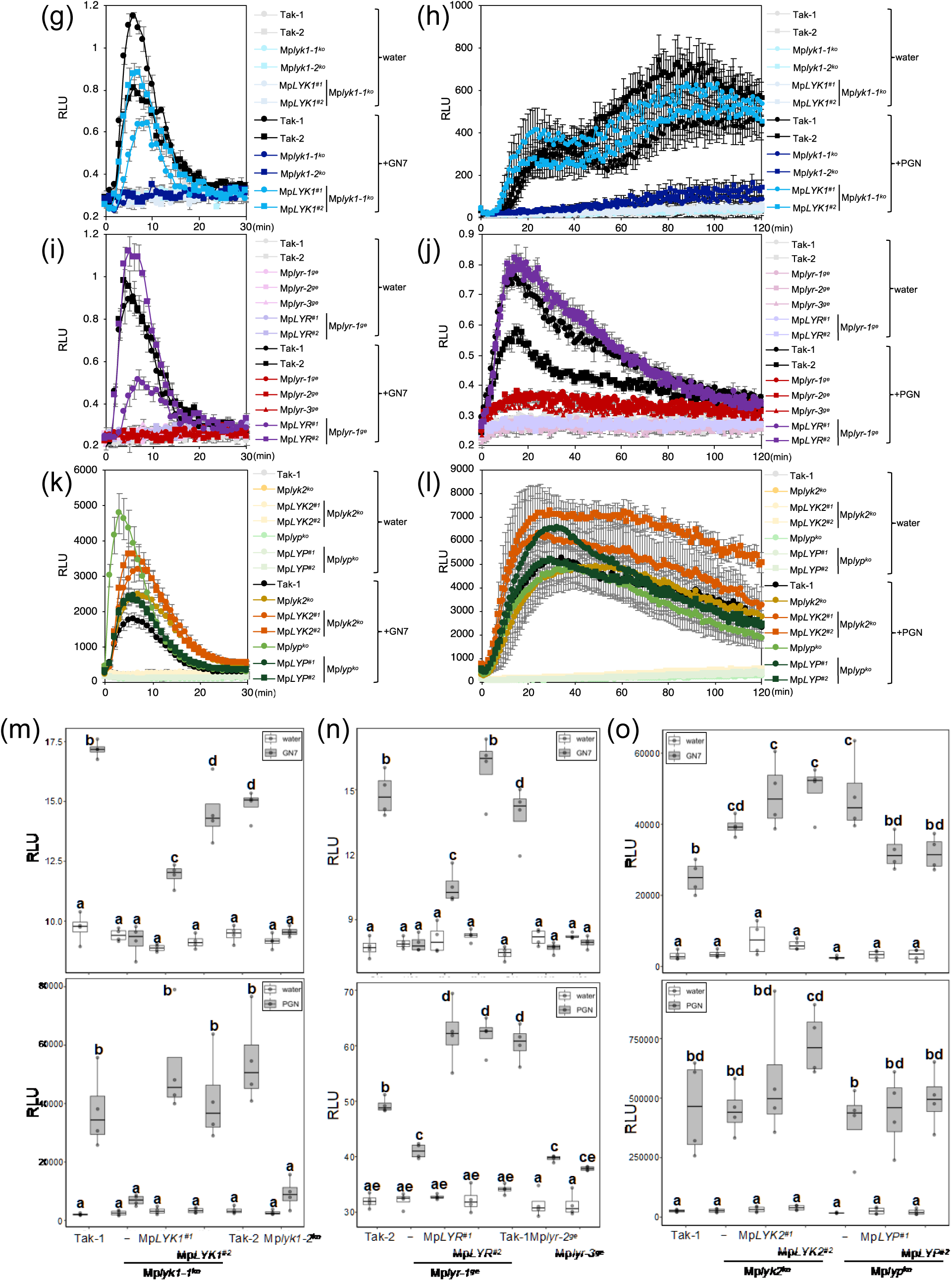
Chitin or PGN-induced ROS burst in LysM receptor homolog disruptants. (a-f) Six-day-old gemmalings of wild-type plants, disruptants, and complementation lines were treated with 1μM GN7 (a, c) or 500 μg/ml PGN from *Bacillus subtilis* (b, d). The boxplot indicates total value of RLU measured by luminometer for 30 minutes after GN7 treatment (e) or for 2 hours after PGN treatment (f). (g-o) Six-day-old gemmalings of independent disruptants or complementation lines were treated with GN7 (g, i, k) or PGN (h, j, l). The boxplot (m-o) indicates total value of RLU measured by luminometer for 30 minutes after GN7 treatment or for 2 hours after PGN treatment. Boxes show upper and lower quartiles of the value, and black lines represent the medians. Statistical groups were determined using the Tukey HSD test. Statistically significant differences are indicated by different letters (p < 0.05).

**Figure S8.**
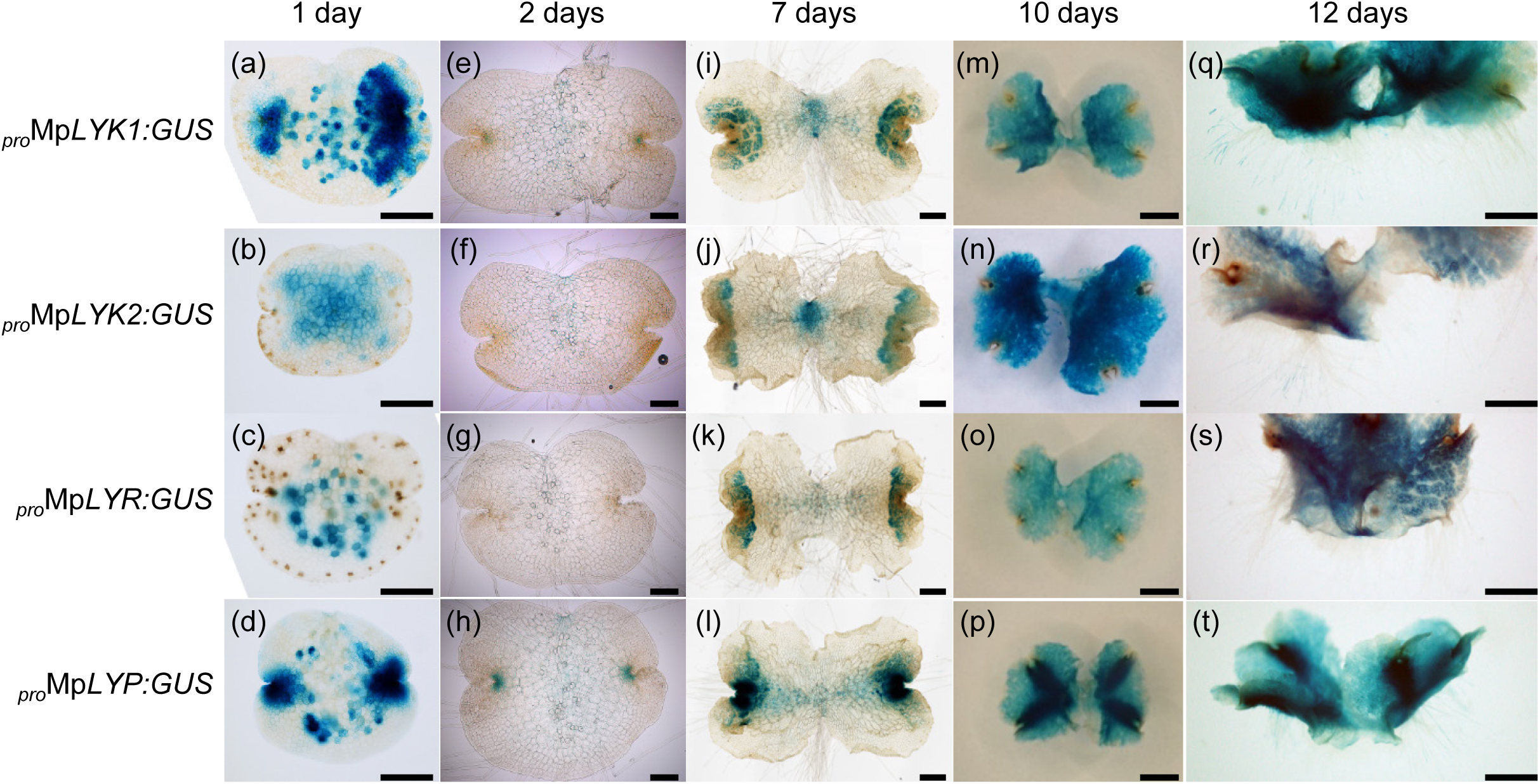
Expression profiles of Mp*LysM* genes. GUS-staining images of plants harboring *_pro_*Mp*LYK1:GUS*, *_pro_*Mp*LYK2:GUS*, *_pro_*Mp*LYR:GUS*, and *_pro_*Mp*LYP:GUS*. (a-d) One-day-old gemmalings. (e-h) Two-day-old gemmalings. (i-l) Seven-day-old gemmalings. (m-p) Ten-day-old thallus, dorsal side. (q-t) 12-day-old thallus with rhizoids. Bars = 200 μm in (a-h), 500 μm in (i-l), and 2 mm in (m-t).

**Figure S9.**
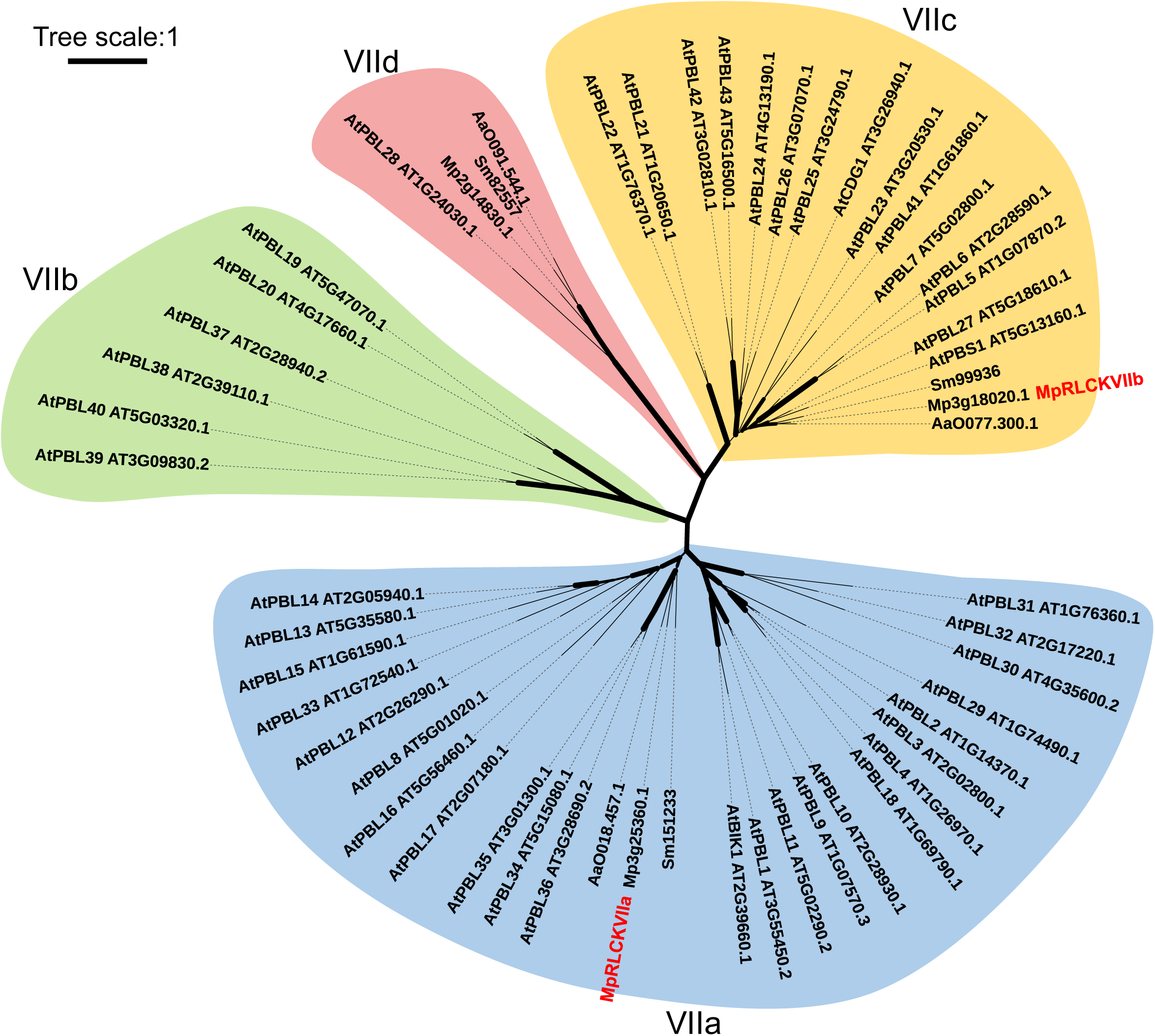
Unrooted phylogenetic tree of the subfamily VII RLCK in plants. The full-length amino acid sequences (Supplementary Table S4) were used for MUSCLE alignment analysis and PhyML tree analysis. A graphical view of the tree has been generated using iTOL. Width of branches denote bootstrap support based on 100 repetitions. The four major subfamilies were designated as VIIa, VIIb, VIIc, and VIId based on the classification by (Shiu et al., 2004).

**Table S1. Amino acid sequences used for drawing phylogenetic tree of LysM proteins**

**Table S2. LysM proteins containing modified P-Loops**

**Table S3. Oligonucleotides used in this study**

**Table S4. Amino acid sequences used for drawing phylogenetic tree of the subfamily VII RLCK**

**Table S5. Phosphoproteome data upon chitin treatment**

**Table S6. Phosphoproteome data upon chitin or blue-light treatment**

**Table S7. RNA-seq data used for Figure 1e**

**Table S8. GO analysis used for Figure 1f**

**Table S9. RNA-seq data used for Supplementary Figure S2a**

**Table S10. GO analysis result used for Supplementary Figure S2b**

**Table S11 RNA-seq data used for Figure 5c**

## References

Bi, G., Zhou, Z., Wang, W., Li, L., Rao, S., Wu, Y., et al. (2018). Receptor-Like Cytoplasmic Kinases Directly Link Diverse Pattern Recognition Receptors to the Activation of Mitogen-Activated Protein Kinase Cascades in Arabidopsis. Plant Cell 30, 1543–1561. doi:10.1105/tpc.17.00981.

Boller, T., and Felix, G. (2009). A renaissance of elicitors: Perception of microbe-associated molecular patterns and danger signals by pattern-recognition receptors. Annu. Rev. Plant Biol. 60, 379–406. doi:10.1146/annurev.arplant.57.032905.105346.

Bowman, J. L., Kohchi, T., Yamato, K. T., Jenkins, J., Shu, S., Ishizaki, K., et al. (2017). Insights into Land Plant Evolution Garnered from the *Marchantia polymorpha* Genome. Cell 171, 287–299.e15. doi:10.1016/j.cell.2017.09.030.

Bressendorff, S., Azevedo, R., Kenchappa, C. S., Ponce de León, I., Olsen, J. V., Rasmussen, M. W., et al. (2016). An innate immunity pathway in the moss *Physcomitrella patens*. Plant Cell 28, 1328–1342. doi:10.1105/tpc.15.00774.

Cao, Y., Liang, Y., Tanaka, K., Nguyen, C. T., Jedrzejczak, R. P., Joachimiak, A., et al. (2014). The kinase LYK5 is a major chitin receptor in *Arabidopsis* and forms a chitin-induced complex with related kinase CERK1. eLife 3, 1504–19. doi:10.7554/eLife.03766.

Carella, P., Gogleva, A., Tomaselli, M., Alfs, C., and Schornack, S. (2018). *Phytophthora palmivora* establishes tissue-specific intracellular infection structures in the earliest divergent land plant lineage. PNAS 115, E3846–E3855. doi:10.1073/pnas.1717900115.

Carlson, M. (2016). KEGG.db: A set of annotation maps for KEGG. R package version 3.2.3. doi:10.18129/B9.bioc.KEGG.db.

Carlson, M. (2019). GO.db: A set of annotation maps describing the entire Gene Ontology. R package version 3.8.2. doi:10.18129/B9.bioc.GO.db.

Carlson, M., Liu, T., Lin, C., Falcon, S., Zhang, J., and MacDonald, J. (2018). PFAM.db: A set of protein ID mappings for PFAM. R package version 3.6.0.

Carotenuto, G., Chabaud, M., Miyata, K., Capozzi, M., Takeda, N., Kaku, H., et al. (2017). The rice LysM receptor like kinase OsCERK1 is required for the perception of short chain chitin oligomers in arbuscular mycorrhizal signaling. New Phytol. 214, 1440–1446. doi:10.1111/nph.14539.

Chen, S., Zhou, Y., Chen, Y., and Gu, J. (2018). fastp: an ultra-fast all-in-one FASTQ preprocessor. Bioinformatics 34, i884–i890. doi:10.1093/bioinformatics/bty560.

Chinchilla, D., Bauer, Z., Regenass, M., Boller, T., and Felix, G. (2006). The *Arabidopsis* Receptor Kinase FLS2 Binds flg22 and Determines the Specificity of Flagellin Perception. Plant Cell 18, 465–476. doi:10.1105/tpc.105.036574.

Chinchilla, D., Zipfel, C., Robatzek, S., Kemmerling, B., Nürnberger, T., Jones, J. D. G., et al. (2007). A flagellin-induced complex of the receptor FLS2 and BAK1 initiates plant defence. Nature 448, 497–501. doi:10.1038/nature05999.

Christie, J. M. (2007). Phototropin Blue-Light Receptors. Annu. Rev. Plant Biol., 21–45. doi:10.1146/annurev.arplant.58.032806.103951.

Cox, J., and Mann, M. (2008). MaxQuant enables high peptide identification rates, individualized p.p.b.-range mass accuracies and proteome-wide protein quantification. Nat. Biotechnol. 26, 1367–1372. doi:10.1038/nbt.1511.

Delaux, P.-M., Radhakrishnan, G. V., Jayaraman, D., Cheema, J., Malbreil, M., Volkening, J. D., et al. (2015). Algal ancestor of land plants was preadapted for symbiosis. PNAS 112, 13390–13395. doi:10.1073/pnas.1515426112.

Dobin, A., Davis, C. A., Schlesinger, F., Drenkow, J., Zaleski, C., Jha, S., et al. (2013). STAR: ultrafast universal RNA-seq aligner. Bioinformatics 29, 15–21. doi:10.1093/bioinformatics/bts635.

Faulkner, C., Petutschnig, E., Benitez-Alfonso, Y., Beck, M., Robatzek, S., Lipka, V., et al. (2013). LYM2-dependent chitin perception limits molecular flux via plasmodesmata. PNAS 110, 9166–9170. doi:10.1073/pnas.1203458110/-/DCSupplemental.

Finn, R. D., Coggill, P., Eberhardt, R. Y., Eddy, S. R., Mistry, J., Mitchell, A. L., et al. (2016). The Pfam protein families database: towards a more sustainable future. Nucleic Acids Res. 44, D279–D285. doi:10.1093/nar/gkv1344.

Gimenez-Ibanez, S., Zamarreño, A. M., García-Mina, J. M., and Solano, R. (2019). An Evolutionarily Ancient Immune System Governs the Interactions between Pseudomonas syringae and an Early-Diverging Land Plant Lineage. Current Biology 29, 2270–2281.e4. doi:10.1016/j.cub.2019.05.079.

Guindon, S., and Gascuel, O. (2003). A Simple, Fast, and Accurate Algorithm to Estimate Large Phylogenies by Maximum Likelihood. Systematic Biology 52, 696–704. doi:10.1080/10635150390235520.

Gust, A. A., Willmann, R., Desaki, Y., Grabherr, H. M., and Nürnberger, T. (2012). Plant LysM proteins: modules mediating symbiosis and immunity. Trends Plant Sci. 17, 495–502. doi:10.1016/j.tplants.2012.04.003.

Humphreys, C. P., Franks, P. J., Rees, M., Bidartondo, M. I., Leake, J. R., and Beerling, D. J. (2010). Mutualistic mycorrhiza-like symbiosis in the most ancient group of land plants. Nat. Commun. 1, 1–7. doi:10.1038/ncomms1105.

Ishizaki, K., Chiyoda, S., Yamato, K. T., and Kohchi, T. (2008). *Agrobacterium*-Mediated Transformation of the Haploid *Liverwort Marchantia* polymorpha L., an Emerging Model for Plant Biology. Plant Cell Physiol. 49, 1084–1091. doi:10.1093/pcp/pcn085.

Ishizaki, K., Johzuka-Hisatomi, Y., Ishida, S., Iida, S., and Kohchi, T. (2013). Homologous recombination-mediated gene targeting in the liverwort *Marchantia polymorpha* L. Sci Rep 3, 1888–6. doi:10.1038/srep01532.

Ishizaki, K., Nishihama, R., Ueda, M., Inoue, K., Ishida, S., Nishimura, Y., et al. (2015). Development of Gateway Binary Vector Series with Four Different Selection Markers for the Liverwort *Marchantia polymorpha*. PLoS ONE 10, e0138876–13. doi:10.1371/journal.pone.0138876.

Ishizaki, K., Nonomura, M., Kato, H., Yamato, K. T., and Kohchi, T. (2012). Visualization of auxin-mediated transcriptional activation using a common auxin-responsive reporter system in the liverwort *Marchantia polymorpha*. J Plant Res 125, 643–651. doi:10.1007/s10265-012-0477-7.

Iwakawa, H., Melkonian, K., Schlüter, T., Jeon, H.-W., Nishihama, R., Motose, H., et al. (2021). *Agrobacterium*-Mediated Transient Transformation of *Marchantia* Liverworts. Plant Cell Physiol. 62, 1718–1727. doi:10.1093/pcp/pcab126.

Jefferson, R. A., Kavanagh, T. A., and Bevan, M. W. (1987). GUSfusions: β-glucuronidaseas a sensitiveandversatilegene fusion marker in higher plants. EMBO J. 6, 3901–3907.

Kadota, Y., Sklenar, J., Derbyshire, P., Stransfeld, L., Asai, S., Ntoukakis, V., et al. (2014). Direct Regulation of the NADPH Oxidase RBOHD by the PRR-Associated Kinase BIK1 during Plant Immunity. Molecular Cell 54, 43–55. doi:10.1016/j.molcel.2014.02.021.

Kaku, H., Nishizawa, Y., Ishii-Minami, N., Akimoto-Tomiyama, C., Dohmae, N., Takio, K., et al. (2006). Plant cells recognize chitin fragments for defense signaling through a plasma membrane receptor. PNAS 103, 11086–11091.

Kanehisa, M., Sato, Y., and Morishima, K. (2016). BlastKOALA and GhostKOALA: KEGG Tools for Functional Characterization of Genome and Metagenome Sequences. Journal of Molecular Biology 428, 726–731. doi:10.1016/j.jmb.2015.11.006.

Kasahara, M., Kagawa, T., Sato, Y., Kiyosue, T., and Wada, M. (2004). Phototropins Mediate Blue and Red Light-Induced Chloroplast Movements in *Physcomitrella patens*. Plant Physiol. 135, 1388–1397. doi:10.1104/pp.104.042705.

Koide, E., Suetsugu, N., Iwano, M., Gotoh, E., Nomura, Y., Stolze, S. C., et al. (2020). Regulation of Photosynthetic Carbohydrate Metabolism by a Raf-Like Kinase in the Liverwort *Marchantia polymorpha*. Plant Cell Physiol. 61, 631–643. doi:10.1093/pcp/pcz232.

Kolde, R. (2019). pheatmap: Pretty Heatmaps. R package version 1.0.12. Available at: https://CRAN.R-project.org/package=pheatmap.

Komatsu, A., Terai, M., Ishizaki, K., Suetsugu, N., Tsuboi, H., Nishihama, R., et al. (2014). Phototropin Encoded by a Single-Copy Gene Mediates Chloroplast Photorelocation Movements in the Liverwort *Marchantia polymorpha*. Plant Physiol., 411–427. doi:10.1104/pp.114.245100.

Kubota, A., Ishizaki, K., Hosaka, M., and Kohchi, T. (2014). *Agrobacterium*-Mediated Transformation of the Liverwort *Marchantia polymorpha* Using Regenerating Thalli. Biosci. Biotechnol. Biochem. 77, 167–172. doi:10.1271/bbb.120700.

Kumar, R., Ichihashi, Y., Kimura, S., Chitwood, D. H., Headland, L. R., Peng, J., et al. (2012). A high-throughput method for Illumina RNA-Seq library preparation. Front. Plant Sci. 3, 1–10. doi:10.3389/fpls.2012.00202/abstract.

Li, F.-W., Nishiyama, T., Waller, M., Frangedakis, E., Keller, J., Li, Z., et al. (2020). *Anthoceros* genomes illuminate the origin of land plants and the unique biology of hornworts. Nat. Plants 6, 259–272. doi:10.1038/s41477-020-0618-2.

Li, J., Wen, J., Lease, K. A., Doke, J. T., Tax, F. E., and Walker, J. C. (2002). BAK1, an *Arabidopsis* LRR Receptor-like Protein Kinase, Interacts with BRI1 and Modulates Brassinosteroid Signaling. Cell 110, 213–222. doi:10.1016/S0092-8674(02)00812-7.

Liang, Y., Cao, Y., Tanaka, K., Thibivilliers, S., Wan, J., Choi, J., et al. (2013). Nonlegumes Respond to Rhizobial Nod Factors by Suppressing the Innate Immune Response. Science 341, 1384–1387. doi:10.1126/science.1242032.

Liu, B., Li, J.-F., Ao, Y., Qu, J., Li, Z., Su, J., et al. (2012). Lysin Motif–Containing Proteins LYP4 and LYP6 Play Dual Roles in Peptidoglycan and Chitin Perception in Rice Innate Immunity. Plant Cell 24, 3406–3419. doi:10.1105/tpc.112.102475.

Lombard, V., Ramulu, H. G., Drula, E., Coutinho, P. M., and Henrissat, B. (2014). The carbohydrate-active enzymes database (CAZy) in 2013. Nucleic Acids Res. 42, D490–D495. doi:10.1093/nar/gkt1178.

Love, M. I., Huber, W., and Anders, S. (2014). Moderated estimation of fold change and dispersion for RNA-seq data with DESeq2. Genome Biol. 15, 1–21. doi:10.1186/s13059-014-0550-8.

Madsen, E. B., Madsen, L. H., Radutoiu, S., Olbryt, M., Rakwalska, M., Szczyglowski, K., et al. (2003). A receptor kinase gene of the LysM type is involved in legume perception of rhizobial signals. Nature 425, 637–640.

Matsumoto, A., Schlüter, T., Melkonian, K., Takeda, A., Nakagami, H., and Mine, A. (2022). A versatile Tn7 transposon-based bioluminescence tagging tool for quantitative and spatial detection of bacteria in plants. Plant Commun. 3, 100227. doi:10.1016/j.xplc.2021.100227.

Miya, A., Albert, P., Shinya, T., Desaki, Y., Ichimura, K., Shirasu, K., et al. (2007). CERK1, a LysM receptor kinase, is essential for chitin elicitor signaling in *Arabidopsis*. PNAS 104, 19613–19618.

Miyata, K., Kozaki, T., Kouzai, Y., Ozawa, K., Ishii, K., Asamizu, E., et al. (2014). The Bifunctional Plant Receptor, OsCERK1, Regulates Both Chitin-Triggered Immunity and Arbuscular Mycorrhizal Symbiosis in Rice. Plant Cell Physiol. 55, 1864–1872. doi:10.1093/pcp/pcu129.

Miyauchi, S., Hage, H., Drula, E., Lesage-Meessen, L., Berrin, J.-G., Navarro, D., et al. (2020). Conserved white-rot enzymatic mechanism for wood decay in the Basidiomycota genus *Pycnoporus*. DNA Res. 27, 1–14. doi:10.1093/dnares/dsaa011.

Miyauchi, S., Navarro, D., Grigoriev, I. V., Lipzen, A., Riley, R., Chevret, D., et al. (2016). Visual Comparative Omics of Fungi for Plant Biomass Deconstruction. Front. Microbiol. 7, 1335. doi:10.3389/fmicb.2016.01335.

Miyauchi, S., Navarro, D., Grisel, S., Chevret, D., Berrin, J.-G., and Rosso, M.-N. (2017). The integrative omics of white-rot fungus *Pycnoporus coccineus* reveals co-regulated CAZymes for orchestrated lignocellulose breakdown. PLoS ONE 12, e0175528. doi:10.1371/journal.pone.0175528.

Miyauchi, S., Rancon, A., Drula, E., Hage, H., Chaduli, D., Favel, A., et al. (2018). Integrative visual omics of the white-rot fungus *Polyporus brumalis* exposes the biotechnological potential of its oxidative enzymes for delignifying raw plant biomass. Biotechnol. Biofuels 11, 1–14. doi:10.1186/s13068-018-1198-5.

Naqvi, S., He, Q., Trusch, F., Qiu, H., Pham, J., Sun, Q., et al. (2022). Blue-light receptor phototropin 1 suppresses immunity to promote *Phytophthora infestans* infection. New Phytol. 233, 2282–2293. doi:10.1111/nph.17929.

Ngou, B. P. M., Heal, R., Wyler, M., Schmid, M. W., and Jones, J. D. G. (2022). Concerted expansion and contraction of immune receptor gene repertoires in plant genomes. Nat. Plants 8, 1146–1152. doi:10.1038/s41477-022-01260-5.

Nishiyama, T., Sakayama, H., de Vries, J., Buschmann, H., Saint-Marcoux, D., Ullrich, K. K., et al. (2018). The Chara Genome: Secondary Complexity and Implications for Plant Terrestrialization. Cell 174, 448–464. doi:10.1016/j.cell.2018.06.033.

Ogata, H., Goto, S., Sato, K., Fujibuchi, W., Bono, H., and Kanehisa, M. (1999). KEGG: Kyoto Encyclopedia of Genes and Genomes. Nucleic Acids Res. 27, 29–34.

Paparella, C., Savatin, D. V., Marti, L., De Lorenzo, G., and Ferrari, S. (2014). The Arabidopsis LYSIN MOTIF-CONTAINING RECEPTOR-LIKE KINASE3 Regulates the Cross Talk between Immunity and Abscisic Acid Responses. Plant Physiol. 165, 262–276. doi:10.1104/pp.113.233759.

R Core Team (2013). A Language and Environment for Statistical Computing. R Foundation for Statistical Computing.

Rawlings, N. D., Barrett, A. J., Thomas, P. D., Huang, X., Bateman, A., and Finn, R. D. (2018). The MEROPS database of proteolytic enzymes, their substrates and inhibitors in 2017 and a comparison with peptidases in the PANTHER database. Nucleic Acids Res. 46, D624– D632. doi:10.1093/nar/gkx1134.

Redkar, A., Ibanez, S. G., Sabale, M., Zechmann, B., Solano, R., and Di Pietro, A. (2022). *Marchantia polymorpha* model reveals conserved infection mechanisms in the vascular wilt fungal pathogen *Fusarium oxysporum*. New Phytol. 234, 227–241. doi:10.1111/nph.17909.

Rensing, S. A., Lang, D., Zimmer, A. D., Terry, A., Salamov, A., Shapiro, H., et al. (2008). The Physcomitrella Genome Reveals Evolutionary Insights into the Conquest of Land by Plants. Science 319, 64–69.

Rich, M. K., Vigneron, N., Libourel, C., Keller, J., Xue, L., Hajheidari, M., et al. (2021). Lipid exchanges drove the evolution of mutualism during plant terrestrialization. Science 372, 864–868. doi:10.1126/science.abg0929.

Roux, M., Schwessinger, B., Albrecht, C., Chinchilla, D., Jones, A., Holton, N., et al. (2011). The Arabidopsis Leucine-Rich Repeat Receptor–Like Kinases BAK1/SERK3 and BKK1/SERK4 Are Required for Innate Immunity to Hemibiotrophic and Biotrophic Pathogens. 23, 2440–2455. doi:10.1105/tpc.111.084301.

Russell, J., and Bulman, S. (2005). The liverwort *Marchantia foliacea* forms a specialized symbiosis with arbuscular mycorrhizal fungi in the genus *Glomus*. New Phytol. 165, 567– 579. doi:10.1111/j.1469-8137.2004.01251.x.

Shimizu, T., Nakano, T., Takamizawa, D., Desaki, Y., Ishii-Minami, N., Nishizawa, Y., et al. (2010). Two LysM receptor molecules, CEBiP and OsCERK1, cooperatively regulate chitin elicitor signaling in rice. Plant J. 64, 204–214. doi:10.1111/j.1365-313X.2010.04324.x.

Shiu, S.-H., Karlowski, W. M., Pan, R., Tzeng, Y.-H., Mayer, K. F. X., and Li, W.-H. (2004). Comparative Analysis of the Receptor-Like Kinase Family in Arabidopsis and Rice. Plant Cell 16, 1220–1234. doi:10.1105/tpc.020834.

Sugano, S. S., Nishihama, R., Shirakawa, M., Takagi, J., Matsuda, Y., Ishida, S., et al. (2018). Efficient CRISPR/Cas9-based genome editing and its application to conditional genetic analysis in *Marchantia polymorpha*. PLoS ONE 13, e0205117–22. doi:10.1371/journal.pone.0205117.

Sugano, S. S., Shirakawa, M., Takagi, J., Matsuda, Y., Shimada, T., Hara-Nishimura, I., et al. (2014). CRISPR/Cas9-Mediated Targeted Mutagenesis in the Liverwort *Marchantia polymorpha* L. Plant Cell Physiol. 55, 475–481. doi:10.1093/pcp/pcu014.

Suzuki, M., Yoshida, I., Suto, K., Desaki, Y., Shibuya, N., and Kaku, H. (2019). AtCERK1 Phosphorylation Site S493 Contributes to the Transphosphorylation of Downstream Components for Chitin-Induced Immune Signaling. Plant Cell Physiol. 60, 1804–1810. doi:10.1093/pcp/pcz096.

Tanaka, K., Nguyen, C. T., Liang, Y., Cao, Y., and Stacey, G. (2013). Role of LysM receptors in chitin-triggered plant innate immunity. Plant Signal. Behav. 8, e22598 147–153. doi:10.4161/psb.22598.

Tatusov, R. L., Fedorova, N. D., Jackson, J. D., Jacobs, A. R., Kiryutin, B., Koonin, E. V., et al. (2003). The COG database: an updated version includes eukaryotes. BMC Bioinform. 4, 1– 14. doi:10.1186/1471-2105-4-41.

The Gene Ontology Consortium (2015). Gene Ontology Consortium: going forward. Nucleic Acids Res. 43, D1049–D1056. doi:10.1093/nar/gku1179.

Thomas, P. D., Campbell, M. J., Kejariwal, A., Mi, H., Karlak, B., Daverman, R., et al. (2003). PANTHER: A Library of Protein Families and Subfamilies Indexed by Function. Genome Research 13, 2129–2141.

Tian, T., Liu, Y., Yan, H., You, Q., Yi, X., Du, Z., et al. (2017). agriGO v2.0: a GO analysis toolkit for the agricultural community, 2017 update. Nucleic Acids Res. 45, W122–W129. doi:10.1093/nar/gkx382.

Tyanova, S., Temu, T., and Cox, J. (2016). The MaxQuant computational platform for mass spectrometry–based shotgun proteomics. Nature Protocols 11, 2301–2319. doi:10.1038/nprot.2016.136.

Willmann, R., Lajunen, H. M., Erbs, G., Newman, M.-A., Kolb, D., Tsuda, K., et al. (2011). Arabidopsis lysin-motif proteins LYM1 LYM3 CERK1 mediate bacterial peptidoglycan sensing and immunity to bacterial infection. PNAS 19824–19829. doi:10.1073/pnas.1112862108/-/DCSupplemental.

Yamada, K., Yamaguchi, K., Shirakawa, T., Nakagami, H., Mine, A., Ishikawa, K., et al. (2016). The *Arabidopsis* CERK1-associated kinase PBL27 connects chitin perception to MAPK activation. EMBO J. 35, 2468–2483. doi:10.15252/embj.201694248.

Zhang, J., Fu, X.-X., Li, R.-Q., Zhao, X., Liu, Y., Li, M.-H., et al. (2020). The hornwort genome and early land plant evolution. Nat. Plants 6, 107–118. doi:10.1038/s41477-019-0588-4.

Zhang, X., Dong, W., Sun, J., Feng, F., Deng, Y., He, Z., et al. (2015). The receptor kinase *CERK1* has dual functions in symbiosis and immunity signalling. Plant J. 81, 258–267. doi:10.1111/tpj.12723.

Zhou, Q., Liu, J., Wang, J., Chen, S., Chen, L., Wang, J., et al. (2020). The juxtamembrane domains of *Arabidopsis* CERK1, BAK1, and FLS2 play a conserved role in chitin induced signaling. J. Integr. Plant Biol. 62, 556–562. doi:10.1111/jipb.12847.

Zhou, Y., Bin Zhou, Pache, L., Chang, M., Khodabakhshi, A. H., Tanaseichuk, O., et al. (2019). Metascape provides a biologist-oriented resource for the analysis of systems-level datasets. Nat. Commun. 10, 1–10. doi:10.1038/s41467-019-09234-6.

Zipfel, C., and Oldroyd, G. E. D. (2017). Plant signalling in symbiosis and immunity. Nature 543, 328–336. doi:10.1038/nature22009.

Zipfel, C., Kunze, G., Chinchilla, D., Caniard, A., Jones, J. D. G., Boller, T., et al. (2006). Perception of the bacterial PAMP EF-Tu by the receptor EFR restricts *Agrobacterium*-mediated transformation. Cell 125, 749–760. doi:10.1016/j.cell.2006.03.037.

